# RUNX2 is a cell-intrinsic brake on Th17 pathogenicity driven by neutrophil extracellular traps

**DOI:** 10.1101/2025.06.26.661866

**Authors:** Caleb Wong Han, Aaron Han Heng, Gabrielle T. Belz, Anne Brüstle, Shaun R. McColl, Iain Comerford

**Affiliations:** Department of Molecular and Biomedical Science, School of Biological Sciences, Faculty of Science Engineering and Technology, The University of Adelaide, Adelaide, SA 5005, Australia; The University of Queensland Frazer Institute, University of Queensland, Brisbane, QLD, 4072, Australia; The John Curtin School of Medical Research, The Australian National University, Canberra, ACT, Australia

**Author notes:** Corresponding author: Dr Iain Comerford.

## Abstract

Th17 cells are crucial mediators of protective immunity against extracellular pathogens but also promote inflammatory responses in pathological settings such as MS. Distinct transcriptional programs can generate Th17 populations termed pathogenic or non-pathogenic. Understanding the transcriptional regulation of these T cell states is emerging and presents opportunities to modulate pathogenic T cell responses. In this study, we demonstrate that pro-inflammatory Th17 cells in MS and EAE express high levels of the transcription factor RUNX2. T cell specific deletion of RUNX2 lowers the threshold for induction of CNS autoimmunity in mice by increasing generation of pathogenic GM-CSF-expressing Th17 cells. RUNX2-dependent inhibition of pathogenic Th17 priming relied on the extent of neutrophil-driven pathogenic Th17 priming *in vivo*. EAE-induced immature neutrophils elicit more pathogenic Th17 cells in the absence of RUNX2 in T cells, an effect reliant on neutrophil-derived extracellular traps. These data uncover a role for RUNX2 in suppressing NET-driven Th17 pathogenicity and suggest that increasing RUNX2 activity in T cells may be a strategy to constrain pathological inflammatory responses where Th17 cells and inflammatory neutrophils interplay.

## Introduction

Th17 cells are particularly implicated in the pathogenesis of pathological tissue inflammation such as the autoimmune neuroinflammatory disease multiple sclerosis (MS), the inflammatory aspects of which are modeled *in vivo* using experimental autoimmune encephalomyelitis (EAE). Th17 cells are observed in active MS lesions in the brain and their presence is correlated with disease severity and relapse ^1^. Supporting their pathogenic role in MS, immunotherapeutics including natalizumab and secukinumab are associated with inhibition of Th17 cell entry into the CNS and their effector cytokines, respectively ^2,3^. In EAE, Th17 cells are required for disease development, and this is elegantly shown in mouse models where deletion of key Th17 transcription factors, such as STAT3, RORγt or IRF4 in T cells renders mice resistant to EAE as a result of limited Th17 responses ^4–6^. However, there is growing evidence that Th17 cells are fundamentally heterogenous and composed of functionally distinct subsets that vary in their inflammatory potential. Understanding these distinctions in Th17 cell biology are important to elucidate their function at mucosal barriers in homeostasis and at effector sites such as the CNS during autoimmune inflammation ^7^. Differentiation of Th17 cells from naïve CD4^+^ T cell precursors is well understood, and their subsequent transition to a pathogenic phenotype from a non-pathogenic phenotype has been demonstrated to require IL-23 signaling, however the underlying transcriptional regulation and network that specifies the pathogenic from non-pathogenic Th17 cell state is not as well defined ^8^. Pathogenic and non-pathogenic Th17 cells can be broadly classified by their production of the proinflammatory cytokine GM-CSF and the anti-inflammatory cytokine IL-10, respectively ^9^. It has been demonstrated that GM-CSF is essential for T cell encephalitogenicity in EAE, which is in contrast to IL-17A and IFNγ that have been shown to be redundant for T cell pathogenicity in EAE ^8^. To date, several transcription factors have been implicated in driving the pathogenic Th17 program, including BHLHE40 and GATA3, which promote T cell expression of GM-CSF ^10,11^. Further, the transcription factors TCF1 and BACH2 are part of the non-pathogenic Th17 transcriptional network, and have recently been shown to limit Th17 cell expansion and pro-inflammatory cytokine production ^12,13^. Critically however, transcriptional repression of the pro-inflammatory program in already established pathogenic Th17 cells has yet to be identified, which constrains development of strategies to reprogram or inhibit pathogenic Th17 responses.

Recent longitudinal analysis of an MS patient cohort revealed that newly developed MS lesions were strongly associated with plasma IL-17A and neutrophil degranulation products ^14^, this suggests that neutrophils may exacerbate the Th17 cell response that contributes to MS lesion formation. There is emerging evidence in experimental models of infection and sterile inflammation that implicate neutrophils in driving Th17 responses ^15–18^, however the specific nature of this interaction in the context of CNS autoimmunity and its regulatory control remains unclear.

In this study, we identified RUNX2 as an inhibitor of Th17 cell pathogenicity in EAE. We found that pathogenic Th17 cells in EAE and MS expressed high levels of RUNX2 and that T cell specific deficiency of *Runx2* lowered the threshold of EAE induction and skewed Th17 cells toward a more pro-inflammatory GM-CSF-secreting phenotype without influencing Treg responses. Moreover, we demonstrate that RUNX2 cell-intrinsically suppresses differentiation of pathogenic Th17 cells driven by neutrophil-derived extracellular DNA. Overall, these results indicate that RUNX2 is expressed by pathogenic Th17 cells and acts as a cell-intrinsic brake on this effector program to limit T cell-driven tissue inflammation in autoimmunity.

## Results

### Pathogenic Th17 cells express high levels of RUNX2

To gain insight into transcriptional regulators that drive dichotomy of Th17 cells during experimental CNS autoimmunity, pathogenic and non-pathogenic Th17 cells were first FACS-sorted for bulk RNA-seq from spleens of EAE-immunized *Il17a-gfp* reporter mice based on reciprocal expression of chemokine receptors CCR2 and CCR6 (Figure. 1A). We have previously shown that a CCR2^+^CCR6^-^ signature enriches for pathogenic GM-CSF-producing Th17 cells, while a CCR2^-^CCR6^+^ signature enriches for non-pathogenic IL-10-producing Th17 cells ^19^. Inspection of the top differentially expressed (DE) genes revealed that CCR2^-^CCR6^+^ non-pathogenic Th17 cells showed enrichment of known ‘stem-like’ Th17 genes ^20^, including *Tcf7*, *Slamf6*, *Sell*, *Ccr7*, *Id3* and *Cd27*, as well as genes encoding the less inflammatory Th17 cytokines, *Il22* and *Il17f*. In contrast, CCR2^+^CCR6^-^ pathogenic Th17 cells exhibited high expression of genes linked with terminal effector, Th1 phenotypes and tissue residency ^20^, including *Nkg7*, *Ifng*, *Tbx21*, *Gzmb*, *Ifitm3* and *Cxcr6* (Figure. 1B). To better understand transcriptional regulation associated with the pathogenic Th17 cell program, we specifically investigated DE genes encoding transcription factors between pathogenic and non-pathogenic Th17 cells (Figure. 1C; Supplementary table 1). A notable transcription factor gene that was strongly DE and significantly upregulated in pathogenic Th17 cells was the gene encoding RUNX2 (Figure. 1C). This stood out as RUNX2 is an important transcription factor in T cell differentiation ^21–24^ but not characterized with respect to its function in pathogenic Th17 cells. To investigate if *RUNX2* was also associated with human pathogenic T cells relevant in the context of MS, we compared transcriptomes of CD4^+^ T cells isolated from the cerebrospinal fluid (CSF) versus the peripheral blood of a cohort of MS patients ^25^. *RUNX2* expression was significantly elevated in CSF-infiltrated CD4^+^ T cells compared to circulating

**Figure 1.**
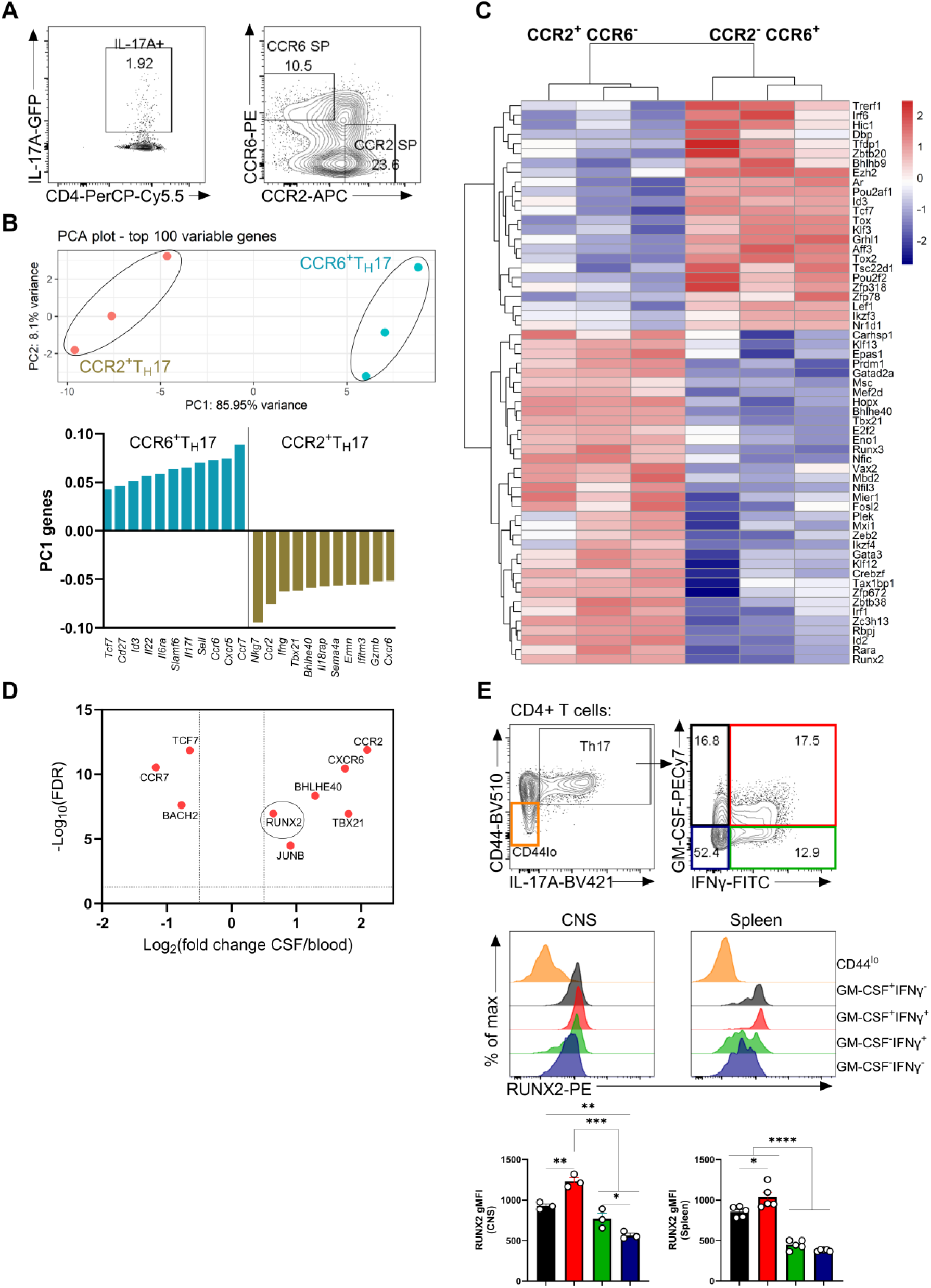
Pathogenic Th17 cells express high levels of RUNX2. **(A)** Gating strategy used to FACS-sort CCR2^+^CCR6^-^ and CCR2^-^CCR6^+^ GFP positive (Th17) cells from the spleens of day 10 EAE-immunized *Il-17a-gfp* mice for RNA-seq. **(B)** Principal component analysis of CCR2^+^CCR6^-^ Th17 and CCR2^-^CCR6^+^ Th17 transcriptomes based on the top 100 variable genes. The top 11 genes, ranked in either the positive or negative direction, representing the most significant sources of variation in the dataset (PC1 axis). **(C)** Heatmap of differentially expressed transcription factors among the significant and differentially expressed genes between CCR2^+^CCR6^-^ and CCR2^-^CCR6^+^ Th17 cells. Heatmap plotted with z scores for each gene calculated using variance stabilized transform values of raw gene count data. Each column represents an individual biological replicate. **(D)** Volcano plot depicting selected genes observed to be enriched in CSF-infiltrating CD4^+^ T cells over peripheral blood-CD4^+^ T cells in MS patients, significance was determined by log2foldchange > ±0.5 and false discovery rate <0.05. **(E)** RUNX2 expression in pathogenic Th17 cells in the CNS and spleen of EAE immunized mice. Representative flow cytometry plots of IL-17A, IFNγ, GM-CSF and RUNX2 expression in CD44^+^CD4^+^ T cells and quantification of RUNX2 geometric MFI in Th17 cell subsets on the basis of their expression of GM-CSF and IFNγ are shown. Each symbol represents a biological replicate, data are shown as mean ± SEM of n=3 mice per group (CNS) and n=5 mice per group (Spleen) from one experiment and 2 independent experiments, respectively. One-way ANOVA with Tukey’s corrections were used for comparisons. *P≤0.05, **P≤0.01 and *** P≤0.001.

CD4^+^ T cells from MS patients, and was part of a program associated with *CXCR6*, *CCR2*, *TXBX21* and *BHLHE40*, indicating that *RUNX2* expression is associated with inflammatory effector CD4^+^ T cells poised to infiltrate the CNS during MS pathogenesis (Figure. 1D). To specifically probe expression of RUNX2 protein in pathogenic Th17 cells, CD4^+^ T cells isolated from the CNS and spleen at the peak of clinical EAE pathogenesis were analyzed. RUNX2 protein was more prominent in pathogenic (GM-CSF^+^IFNγ^-^ and GM-CSF^+^IFNγ^+^) Th17 cell subsets than in Th17 cells that did not express GM-CSF (GM-CSF^-^IFNγ^+^ and GM-CSF^-^IFNγ^-^). Additionally, IL-17A^+^GM-CSF^+^IFNγ^+^ Th17 cells displayed a significant increase in RUNX2 expression compared to IL-17A^+^GM-CSF^+^IFNγ^-^ Th17 cells indicating that RUNX2 expression correlates with Th17 cells expressing the greatest number of pro-inflammatory cytokines (Figure. 1E).

### Regulation of Runx2 expression by Th17-specifying transcription factors

To understand the molecular basis of *Runx2* expression in Th17 cells, regulation of *Runx2* expression by established Th17 transcription factors (RORγt, STAT3, IRF4, JunB, BATF, c-Maf, HIF1α, FOSL2, IKZF3, and TCF1) was evaluated. RNA-seq comparing Th17 polarised cell cultures from transcription factor knockout naïve CD4^+^ T cells with wild-type naïve CD4^+^ T cells (GSE40918, GSE98413) and RNA-seq comparing *Tcf7*-deficient and sufficient Th17 cells from EAE immunized mice (GSE233908) were analysed. Notably, deletion of *Junb* or *Batf* significantly reduced *Runx2* expression in Th17 cells compared to WT, while deletion of *Tcf7* significantly increased *Runx2* expression in Th17 cells compared to WT. However, *Runx2* transcripts were not significantly altered by deletion of *Rorc, Stat3, Irf4, Maf, Hif1a, Fosl2 or Ikzf3* (Figure. 2A). Next, to determine if JunB or BATF bound to proximal putative enhancers upstream of the *Runx2* gene, Th17 cell ATAC-seq and ChIP-seq (SRX15818878, SRX187191, SRX2773920, and SRX187257) were analysed. Notably, a putative proximal enhancer element upstream of the *Runx2* gene marked by overlapping ATAC-seq and histone acetyl transferase EP300 peaks was identified, and in this region, there were clearly detectable JunB and BATF CHIP-seq peaks (Figure. 2B). At this putative enhancer element, JunB and BATF may co-localize with the EP300 transcriptional co-activator to directly activate RUNX2 expression in Th17 cells. RNA-seq of a time-course of Th17 cell polarisation from naïve CD4^+^ T cells (GSE206304) was analysed to investigate whether *Runx2* expression early after TCR-engagement correlates with the upregulation of *Batf* and *Junb*, and the downregulation of *Tcf7* (Figure. 2C). *Runx2* expression was significantly induced in the first 12 hours of culture and induced again between 20–48 hours of culture. In contrast, *Batf* expression was significantly induced between 1–6 hours but decreased between 20–48 hours. *Junb*, which was highly expressed in naïve cells, was significantly reduced between 1–6 hours and increased between 20–48 hours. Finally, *Tcf7* expression was significantly reduced between 1–6 hours and decreased further between 20–48 hours (Figure. 2C). Collectively, these data support a model whereby *Runx2* is positively and directly regulated by BATF and JunB in Th17 cells, with BATF required for early *Runx2* induction and JunB solidifying later *Runx2* expression. Additionally, the data indicate that *Runx2* in Th17 cells is negatively regulated by TCF1.

**Figure 2.**
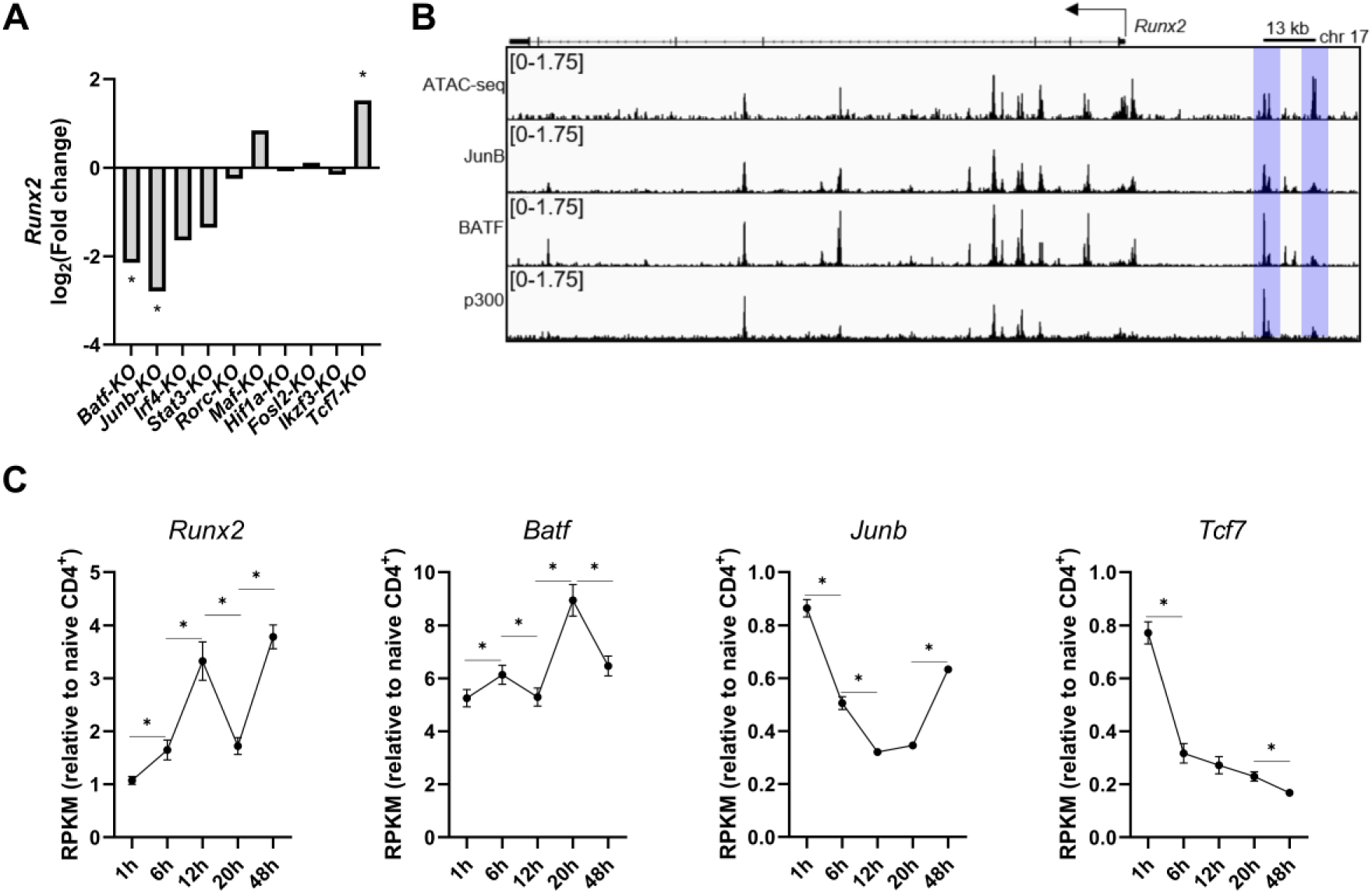
Regulation of *Runx2* expression in Th17 cells by JunB, BATF and TCF1. **(A)** Analysis of *Runx2* expression in various transcription factor knockout Th17 cultures relative to wild-type Th17 cultures after 48h of polarization, and in TCF1 knockout Th17 relative to wild-type Th17 isolated from mice. **(B)** Analysis of JunB, BATF and p300 ChIP-seq tracks, and ATAC-seq data of the *Runx2* locus in Th17 cells after 48h polarization. **(C)** Analysis of *Runx2, Batf, Junb* and *Tcf7* expression in Th17 cultures at 1, 6, 12, 20 and 48h post polarization. Differential analysis was performed using DESeq2 or edgeR, all adjusted P values shown are *P≤2E^-3^.

### T cell expressed RUNX2 inhibits generation of pathogenic Th17 cells

To investigate the functional role of RUNX2 in pathogenic Th17 cell biology, a post-thymic T cell specific *Runx2* knockout mouse strain was developed. T cell specific *Runx2* knockout mice, henceforth referred to as *Runx2^f/f^dLckCre^+^* mice, were generated by crossing *Runx2^fl/fl^* mice ^26^ to *dLck-hCre^3779^*mice ^27,28^, and the conditional deletion of RUNX2 in T cells of *Runx2^f/f^dLckCre^+^*mice was validated by flow cytometry (Figure. S1). We first assessed the T and B lymphocyte compartments in these mice in secondary lymphoid organs. We detected comparable frequencies of T and B cells in the lymph nodes between *Runx2^f/f^dLckCre^+^* mice and *Runx2^fl/fl^* littermate controls, and subset analysis of T cells revealed no differences in the frequencies of effector memory, central memory, or naïve cells in both CD4^+^ and CD8^+^ T cell compartments. In spleen, no differences in the frequency of T or B cells were detected, and the frequencies of central memory and naïve CD4^+^ and CD8^+^ T cells were also comparable between *Runx2^f/f^dLckCre^+^*and *Runx2^fl/fl^* littermate controls. The only statistically significant difference caused by deletion of *Runx2* in the T cell compartment that was detected at homeostasis was a small increase in effector memory cells (between 1%-4%) in both CD4^+^ and CD8^+^ T cell compartments of *Runx2^f/f^dLckCre^+^*compared to littermate controls in the spleen (Figure. S2). Thus, deletion of RUNX2 in post-thymic T cells does not overtly alter the composition of T cells at homeostasis.

Next, the effect of mature T cell restricted *Runx2* deletion on EAE development was assessed using two complementary EAE models, which differ in production of pathogenic Th17 cells and which we refer to hereafter as low adjuvant (LA)-EAE and high adjuvant (HA)-EAE. We used these two models as they differ markedly in the production of pathogenic Th17 cells. The LA-EAE and HA-EAE models involve MOG_35-55_ peptide immunizations in complete Freund’s adjuvant containing either 50µg or 415µg of desiccated *Mycobacterium tuberculosis* H37Rα per mouse, respectively. Compared with HA-EAE immunized mice, LA-EAE mice develop more limited EAE clinical disease severity, generate fewer pathogenic Th17 cells in spleen during priming and fewer pathogenic Th17 cells are recruited to the CNS at peak disease (Figure. S3). The exacerbation in clinical EAE scores and pathogenic Th17 responses in HA-EAE immunized mice was not a consequence of blunted regulatory T cell responses, as analysis of Treg cells revealed comparable frequencies and absolute numbers in the CNS at peak disease in HA-EAE immunized mice relative to LA-EAE immunized mice (Figure. S4). These two models therefore allowed for the effect of RUNX2 deletion under both robust and limited pathogenic Th17 priming conditions *in vivo* to be assessed. *Runx2^f/f^dLckCre^+^* mice and wildtype littermate controls were immunized for either HA-EAE or LA-EAE. Notably, in the LA-EAE model *Runx2^f/f^dLckCre^+^* mice exhibited exacerbated EAE severity, incidence and onset relative to controls, and no difference in disease severity were noted between these genotypes in HA-EAE (Figure. 3A-B; Figure. S6A; Supplementary table 2). These data indicated that deletion of RUNX2 in T cells lowers the threshold for induction of EAE and limited EAE pathogenesis.

**Figure 3.**
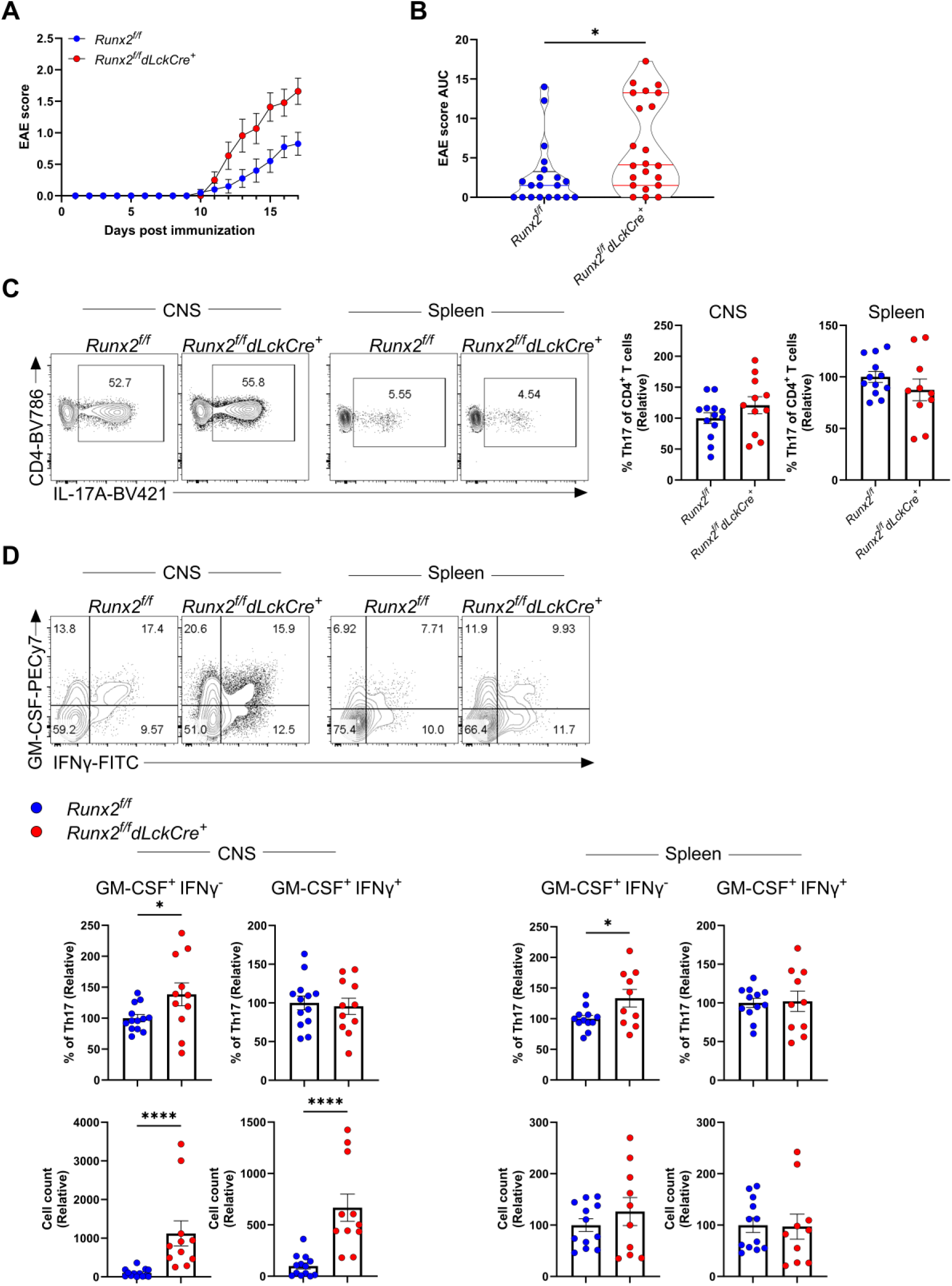
T cell deletion of RUNX2 lowers the threshold for EAE induction and increases pathogenic Th17 generation. **(A)** EAE scores and **(B)** AUC analysis of *Runx2^f/f^* and *Runx2^f/f^dLckCre^+^*mice immunized for LA-EAE. Shown is mean ± SEM of n=20 (*Runx2^f/f^*) and n=22 (*Runx2^f/f^dLckCre^+^*) across 5 independent experiments. Mann-Whitney tests were used for comparisons of AUC. *P≤0.05. **(C)** Representative flow cytometry plots of Th17 from the CNS and spleen on day 13. Percentage of Th17 cells was calculated and normalised to the *Runx2*^f/f^ group in each experiment. **(D)** Representative flow cytometry plots of GM-CSF and IFNγ expression in Th17 cells in the CNS and spleen, the percentage and number of GM-CSF^+^IFNγ^-^ and GM-CSF^+^IFNγ^+^ cells was calculated, normalised to the *Runx2*^f/f^ group in each experiment. Each symbol represents a biological replicate, data are shown as mean ± SEM of n=12-13 (*Runx2*^f/f^) and n=10-11 (*Runx2*^f/f^*dLckCre*^+^) mice across 4 independent experiments. Unpaired t-tests were used for comparisons. *P≤0.05, **P≤0.01 and *** P≤0.001.

To investigate the cellular basis of the RUNX2-dependent disease exacerbation phenotype observed in the LA-EAE model, CD4^+^ T cells isolated from the CNS and spleen of *Runx2^f/f^dLckCre^+^*and control littermate mice at peak EAE disease were analysed. No significant differences in the frequencies of Th17 cells as a proportion of CD4^+^ T cells were apparent between control and *Runx2^f/f^dLckCre^+^* mice (Figure. 3C). However, analysis of CNS-infiltrated Th17 revealed that *Runx2^f/f^dLckCre^+^* mice displayed a significant increase in GM-CSF co-expression compared to littermate controls, in particular in the GM-CSF^+^IFNγ^-^ subset (Figure. 3D). The frequency of GM-CSF^+^IFNγ^-^ Th17 cells was also elevated in the spleens of *Runx2^f/f^dLckCre^+^* mice compared to littermate controls (Figure. 3D), collectively indicating a skewing of Th17 cells to a predominantly pathogenic phenotype at sites of priming and effector function in the absence of RUNX2 in T cells.

To probe whether the RUNX2-dependent disease exacerbation in the LA-EAE model could be explained by attenuated regulatory T cell responses, we analysed Treg cells in the CNS and spleen. In *Runx2^f/f^dLckCre^+^*mice, significant increases in the frequency and number of Treg cells in the CNS were apparent compared to control littermates, but no differences in their quantities in the spleen were measured. This indicates that RUNX2 does not affect Treg cell generation in EAE and it is likely that increased Tregs in the CNS in *Runx2^f/f^dLckCre^+^* mice is due to the increased inflammatory responses ongoing in the CNS of these mice (Figure. S5). Overall, these data do not indicate that impaired Treg responses explain the increase in disease severity seen in the LA-EAE model in *Runx2^f/f^dLckCre^+^* mice. In line with the similar extent of EAE severity in *Runx2^f/f^dLckCre^+^* and control littermate mice in the HA-EAE model, there were no differences in the frequencies of Th17 and Treg cells and the proportions or absolute numbers of GM-CSF-co-expressing Th17 subsets in the CNS and spleen in the HA-EAE model (Figure. S6B-C). Collectively these data indicate that RUNX2 constrains pathogenic Th17 cell generation in the LA-EAE model, a setting where pathogenic Th17 cell production is sub-maximal.

### RUNX2 suppresses pathogenic Th17 cell generation in response to signals from neutrophils

The data thus far suggested that RUNX2 inhibited GM-CSF-co-expressing Th17 cells when adjuvant-induced activation of innate immunity during EAE immunization was limited. We therefore reasoned that RUNX2 did not have an essential role in suppressing pathogenic Th17 cell production in general. In line with this, we found no role for RUNX2 in generation of either Th17 cells generally or pathogenic GM-CSF-secreting Th17 cells following *in vitro* culture of naïve CD4^+^ T cells to Th17 cells using polarising cytokines and CD3/CD28 engaging antibodies (Figure. 4A). These findings suggested that *in vivo* RUNX2 may exert its effects on pathogenic Th17 cell generation by integrating signals arising from cross-talk with the innate immune system that differ between the LA-EAE and HA-EAE models. To investigate the nature of the distinct *in vivo* settings encountered by T cells in these models, we compared the cellular immune responses in the spleens of EAE mice (Figure. 4B; Figure. S7). Analyses of neutrophils, dendritic cells, NK cells, macrophages, monocytes, T cells and B cells were performed at day 11 of EAE, prior to clinical disease onset (Figure. 4B). Comparable numbers of CD45^+^ cells were observed in the spleen in both settings (Figure. 4B) but analysis of the cellular composition of this compartment revealed a significant increase only in the numbers of neutrophils in HA-EAE immunized mice compared to LA-EAE, while no significant differences in dendritic cells, monocytes, NK cells, B cells or T cells were observed (Figure. 4B). Given this stark effect on neutrophil numbers in the spleen driven by adjuvant during the T cell priming phase of EAE and previous reports that neutrophil depletion leads to attenuated EAE severity ^29,30^, we reasoned that neutrophils might promote pathogenic Th17 cell priming. We first utilized an *in vitro* neutrophil and CD4^+^ T cell co-culture system to investigate this. Naïve CD4^+^ T cells were co-cultured with neutrophils isolated from the spleens of EAE mice at different neutrophil:CD4^+^ T cell ratios under Th17 polarizing conditions for 72 hours and analysed by flow cytometry. Analysis of Th17 frequencies at the various neutrophil to CD4^+^ T cell ratios revealed that neutrophils from EAE mice directly promoted Th17 cell priming (Figure. 4C). Further, these neutrophils drove pathogenic Th17 responses as increased ratios of GM-CSF-secreting to IL-10-secreting Th17 cells were apparent with increasing frequencies of neutrophils in the co-cultures (Figure. 4D). We next tested whether RUNX2 in T cells modulated pathogenic Th17 cell induction driven by EAE-derived neutrophils. To test this, we utilised neutrophil:CD4^+^ T cell co-cultures using splenic neutrophils from EAE mice and naïve CD4^+^ T cells from *Runx2^f/f^dLckCre^+^* and control mice. Notably, when EAE-derived neutrophils were added, RUNX2-deficient CD4^+^ T cells demonstrated greater ratios of GM-CSF- to IL-10-expressing Th17 cells compared to control CD4^+^ T cells, indicating that RUNX2 suppresses pathogenic Th17 skewing driven by neutrophils (Figure. 4E).

**Figure 4.**
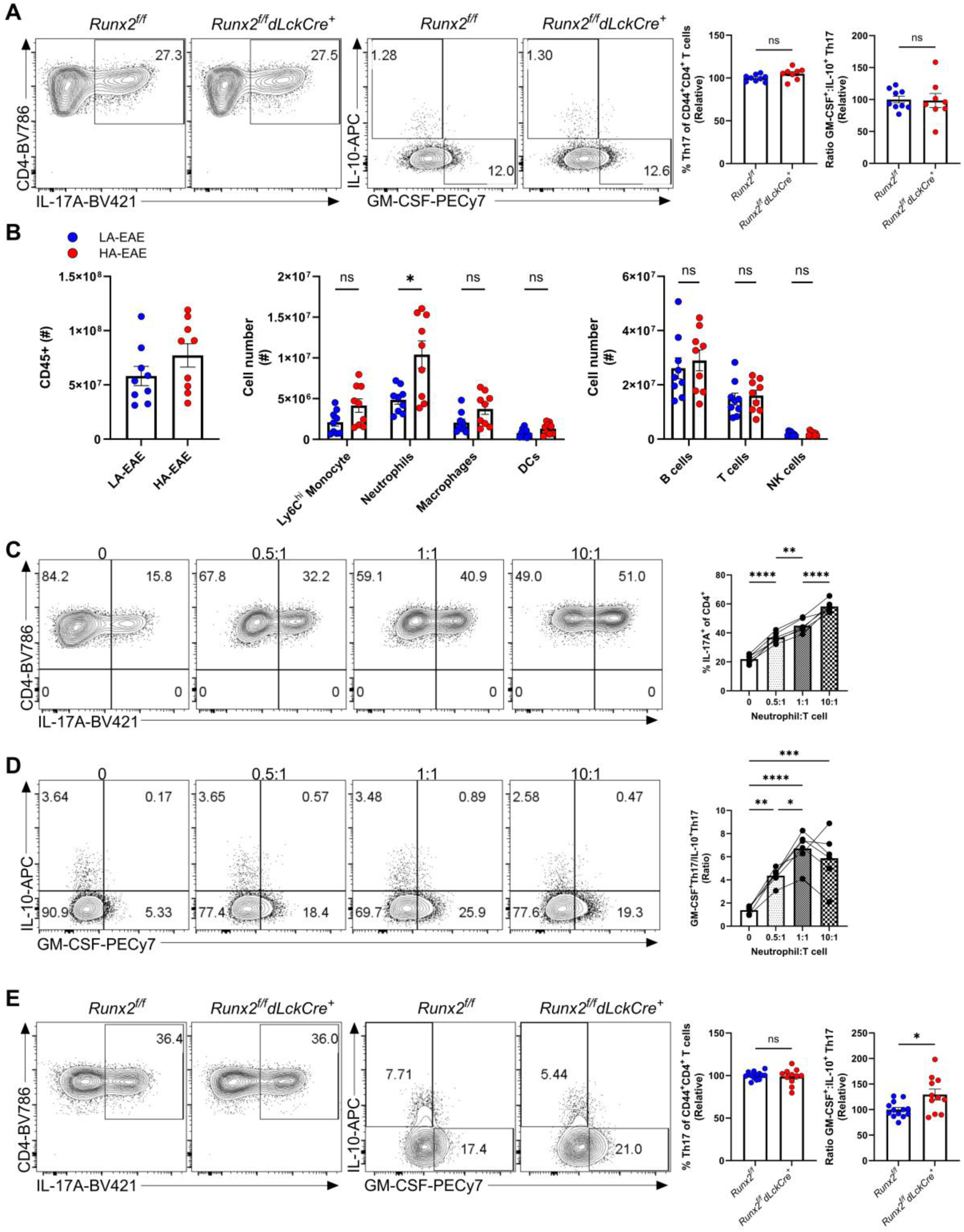
RUNX2 suppresses neutrophil-driven pathogenic Th17 cell generation. **(A)** Naïve CD4^+^ T cells isolated from the spleens of *Runx2^f/f^*and *Runx2^f/f^dLckCre^+^* mice and activated under Th17 polarizing conditions for 72h. Representative flow cytometry plots of IL-17A expression in CD44^+^CD4^+^ T cells and of IL-10 and GM-CSF expression in Th17 cells. Quantification of Th17 and GM-CSF^+^ Th17 to IL-10^+^ Th17 cell ratio normalised to the *Runx2^f/f^* group. **(B)** Quantification of total CD45^+^ cells, myeloid and lymphocyte subsets in the spleen on day 11 following immunisation for HA-EAE or LA-EAE. **(C)** Representative flow cytometry plots of IL-17A expression on CD4^+^ T cells and quantification of the frequency of Th17 cells cultured at various neutrophil ratios. **(D)** Representative flow cytometry plots of GM-CSF and IL-10 expression in Th17 cells and ratios of pathogenic (GM-CSF^+^IL-10^-^) to non-pathogenic (IL-10^+^GM-CSF^-^) Th17 cells. **(E)** Naïve CD4^+^ T cells isolated from *Runx2^f/f^* or *Runx2^f/f^dLckCre^+^* mice cultured with EAE-induced neutrophils at a ratio of 0.5:1 neutrophils to CD4^+^ T cells for 72h under Th17 polarizing conditions. Representative flow cytometry plots of IL-17A expression in CD44^+^CD4^+^ T cells and of IL-10 and GM-CSF expression in Th17 cells. Quantification of Th17 and GM-CSF^+^ Th17 to IL-10^+^ Th17 cell ratio normalised to the *Runx2^f/f^* group. **(A)** Data are mean ± SEM of n=9 (*Runx2^f/f^*) and n=8 (*Runx2^f/f^dLckCre^+^*) mice across three independent experiments. **(B)** Each symbol represents a biological replicate. Data are mean ± SEM of n=9 mice per group across 2 independent experiments. Mann-Whitney tests were used for comparisons. **(C-D)** Each symbol represents a biological replicate. Data are mean ± SEM of n=6 mice per group across 2 independent experiments and normalised to the ‘0:1 neutrophil:CD4^+^ T cell’ group in each experiment. For comparisons, matched repeated measures one-way ANOVA with Tukey corrections was utilised. **(E)** Data are mean ± SEM of n=13 (*Runx2^f/f^*) and n=11 (*Runx2^f/f^dLckCre^+^*) mice across four independent experiments. Each symbol represents a biological replicate. Unpaired t-tests were used for comparisons. *P≤0.05, **P≤0.01 and *** P≤0.001.

Next, we queried whether neutrophils drive pathogenic Th17 cells *in vivo* and whether the RUNX2-dependent increase in EAE pathogenicity was neutrophil-dependent. To test this, mice were immunized for HA-EAE and treated with either anti-Ly6G antibodies to deplete neutrophils, or an isotype control antibody (Fig. 5A). Assessment of neutrophil depletion by intracellular Ly6G staining was performed to overcome surface Ly6G unavailability as previously described ^31^. As expected, anti-Ly6G-treated mice had markedly reduced circulating neutrophils and any remaining neutrophils exhibited reduced Ly6G expression indicating that these were recently generated and not “escapees” of anti-Ly6G treatment (Figure. S8). Strikingly, neutrophil-depleted mice displayed significant reductions in the frequency and numbers of GM-CSF-co-expressing pathogenic Th17 cells in spleen on day 11 of EAE, indicating that neutrophils promote pathogenic Th17 cell priming in EAE (Figure. 5B). Neutrophil depletion did not affect the magnitude of Treg responses in the spleen (Figure. S9). Next, we questioned whether the increase in EAE clinical severity observed in the LA-EAE model in *Runx2^f/f^dLckCre^+^* mice compared to wildtype littermates controls was dependent on neutrophils. Unlike isotype control treated *Runx2*^f/f^*dLckCre*^+^ mice, neutrophil depleted *Runx2*^f/f^*dLckCre*^+^ mice did not display increased clinical EAE compared to isotype control treated wildtype littermates (Figure. 5C), indicating that the increased EAE severity in *Runx2*^f/f^*dLckCre*^+^ mice is neutrophil dependent. We next tested if RUNX2 expression in Th17 cells varied with respect to neutrophil-derived signals. Increased RUNX2 expression was apparent in Th17 cells induced in HA-EAE immunized mice compared to LA-EAE immunized mice (Figure. S10A), and increased RUNX2 expression in Th17 cells in a co-culture system correlated with increasing frequencies of neutrophils (Figure. S10B). Together, these data collectively demonstrate that RUNX2 modulates crosstalk between neutrophils and CD4^+^ T cells during EAE priming that drives pathogenic Th17 cells.

**Figure 5.**
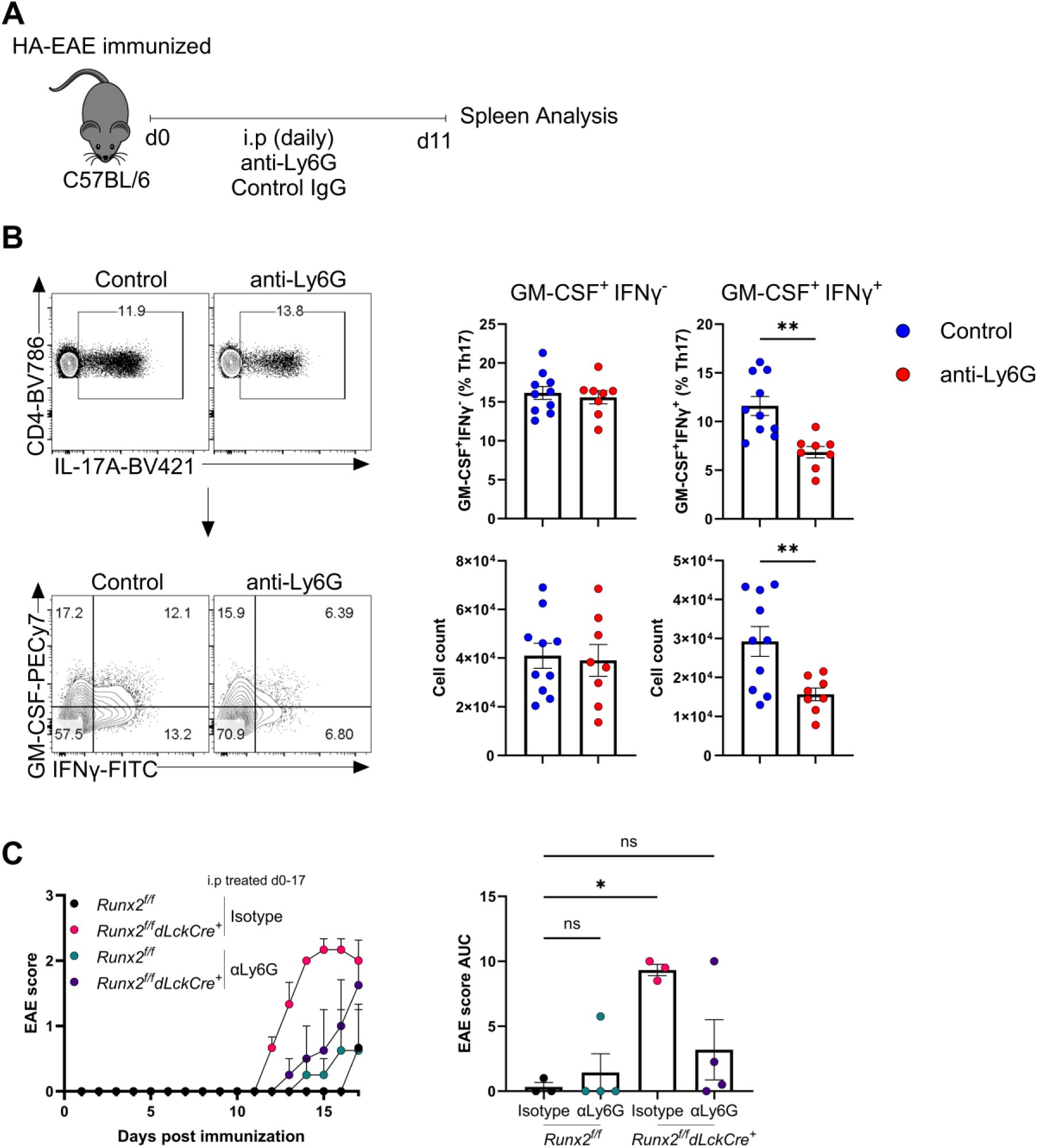
Inhibition of EAE by RUNX2 expression in T cells is dependent on neutrophils. **(A)** Schematic of neutrophil depletion in HA-EAE immunized mice. Mice were IP injected with anti-Ly6G or isotype control antibodies daily and humanely euthanised before disease onset (day 11). **(B)** Representative flow cytometry plots of GM-CSF and IFNγ expression in Th17 cells from the spleen (pre-onset, day 11) and quantification of GM-CSF^+^IFNγ^-^ and GM-CSF^+^IFNγ^+^ Th17 cells. **(C)** *Runx2^f/f^* or *Runx2^f/f^dLckCre^+^*mice immunized for LA-EAE treated with isotype control or anti-Ly6G antibodies. EAE scores and AUC analysis. Shown is mean ± SEM of n=3 (isotype-*Runx2^f/f^*), n=3 (isotype-*Runx2^f/f^dLckCre^+^*), n=4 (anti-Ly6G-*Runx2^f/f^*) and n=4 (anti-Ly6G-*Runx2^f/f^dLckCre^+^*) from one experiment. Ordinary one-way ANOVA with Dunnett’s corrections were used for comparisons of AUC. *P≤0.05. **(A-B)** Each symbol represents a biological replicate, data are mean ± SEM of n=10 (isotype) and n=8 (anti-Ly6G) across 2 independent experiments. Unpaired t-tests were used for comparisons. *P≤0.05, **P≤0.01 and *** P≤0.001.

### Extracellular DNA released from immature neutrophils drives pathogenic Th17 cells in the absence of RUNX2

To gain further insight, we probed the nature of the crosstalk between neutrophils and T cells during EAE that is modulated by RUNX2 in more detail. First, we determined the subset of neutrophil responsible for driving pathogenic Th17 cells in EAE. Neutrophil maturity and immaturity were determined by surface expression of markers CXCR2 and CD49d, respectively, as previously described ^32,33^. To confirm whether this pattern was consistent in the EAE model, we evaluated Ly6G expression levels in CXCR2- and/or CD49d-expressing neutrophil subsets from HA-EAE and LA-EAE immunized mice, as Ly6G expression is positively correlated with neutrophil maturity ^34^. Across both EAE immunization types, CXCR2^+^CD49d^−^ neutrophils exhibited higher Ly6G expression, while CXCR2^-^CD49d^+^ neutrophils showed lower Ly6G expression, consistent with previous findings. This supports the use of this strategy to distinguish mature and immature subsets in the EAE model. Notably, we also observed a CXCR2^−^CD49d^−^ neutrophil population with intermediate Ly6G expression, potentially representing a transitional neutrophil subset progressing toward maturity (Figure. 6A). In the spleen on day 11 post-immunisation, an increase in the absolute numbers of CXCR2^+^CD49d^-^ (mature), CXCR2^-^CD49d^+^ (immature) and CXCR2^-^CD49d^-^ (intermediate) neutrophils was apparent in HA-EAE compared to LA-EAE. However, amongst splenic neutrophils, a significantly higher percentage were of an immature phenotype in the spleens of HA-EAE immunized compared to LA-EAE immunized mice (Figure. 6A). These data indicate that there is substantial mobilisation of immature neutrophils to the spleen driven by adjuvant-induced signals during immune priming in EAE. We reasoned that these immature neutrophils may be responsible for driving increased pathogenic Th17 cells. To test this, we enriched total neutrophils from spleens of EAE mice and also further FACS-sorted immature, mature and intermediate neutrophils for co-culture with naïve CD4^+^ T cells during polarisation to a Th17 phenotype. Strikingly, the capacity to promote pathogenic Th17 cells was confined to immature neutrophils (Figure. 6B). Next, we queried the mechanism behind pathogenic Th17 cell polarisation this crosstalk. Previous work in models of infection and sterile inflammation have shown that neutrophils can drive Th17 differentiation through their release of granule proteins, and the release of extracellular DNA complexed with histones in neutrophil extracellular traps (NETs) ^15–18^. Therefore, we tested whether histones were required for the crosstalk between EAE-derived neutrophils and pathogenic Th17 cells using *Tlr2^-/-^* CD4^+^ cells defective in detection of extracellular histones ^18,35^. In the presence of neutrophils, no differences between control and *Tlr2^-/-^* T cells were apparent with respect to general Th17 and pathogenic Th17 cell polarisation (Figure. S11). Therefore, we next examined the contribution of extracellular DNA. Recent studies in the EAE model have demonstrated that NETs contribute to increased CNS inflammatory responses, and their depletion by *in vivo* administered Dnase1 reduces disease severity and decreases CNS-infiltrating Th17 cells ^36^. To test whether RUNX2 modulates neutrophil-driven pathogenic Th17 responses driven by NETs, naïve CD4^+^ T cells were isolated from *Runx2*^f/f^*dLckCre*^+^ and control mice and co-cultured with neutrophils with or without Dnase1 treatment for 72 hours under pathogenic Th17 cell-polarizing conditions (Figure. 6C). Strikingly, the increase in pathogenic Th17 skewing observed in RUNX2-deficient CD4^+^ T cells cultured with neutrophils was abolished by Dnase1 treatment, while no differences were observed in control CD4^+^ T cells, demonstrating that extracellular DNA is an essential component of the neutrophil derived signal that promotes pathogenic Th17 cells that is modulated by RUNX2 (Figure. 6C). Collectively these data establish RUNX2 as a cell intrinsic brake expressed by pathogenic Th17 cells that restrains pathogenic effector functions driven by detection of NETs.

**Figure 6.**
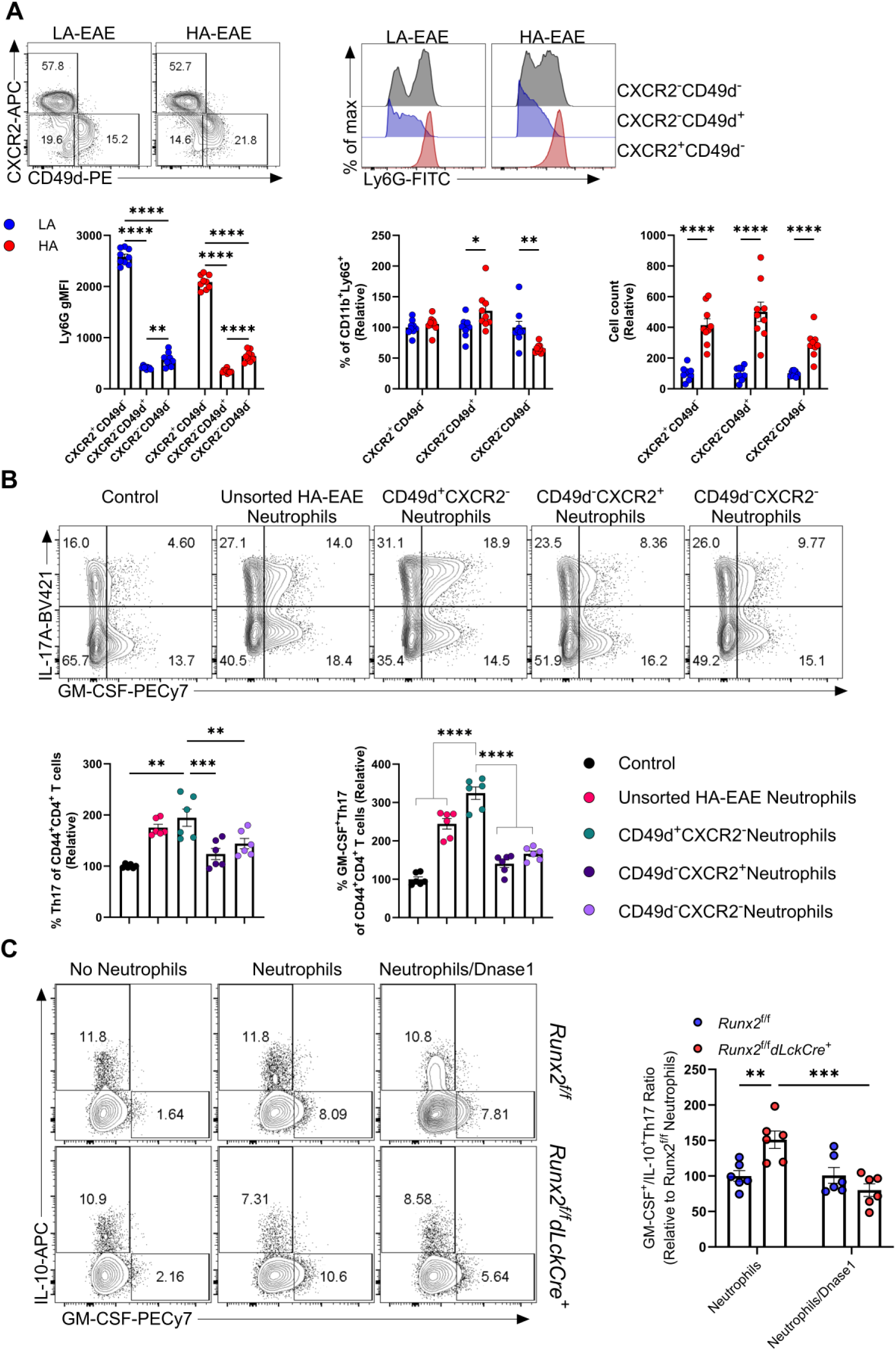
RUNX2 limits pathogenic Th17 cells induced by immature neutrophil-derived extracellular DNA. **(A)** Representative analysis of Ly6G, CXCR2 and CD49d expression on neutrophils in the spleen on day 11. Ly6G geometric MFI of CXCR2^+^CD49d^-^ and CXCR2^-^CD49d^+^ and CXCR2^-^ CD49d^-^ neutrophils, and the percentages and numbers of each neutrophil subset was calculated. Percentages and numbers normalised to the LA-EAE group in each experiment. **(B)** Naïve CD4^+^ T cells cultured under Th17 polarizing conditions in the absence (Control) or presence of neutrophils (0.5:1 neutrophil:CD4^+^ T cell ratio). Neutrophils either enriched from spleen of d11 HA-EAE-immunized mice (Unsorted HA-EAE neutrophils) or FACS-sorted on CXCR2 and/or CD49d. Representative flow cytometry plots of IL-17A and GM-CSF expression in CD44^+^CD4^+^ T cells. Quantification of Th17 and GM-CSF^+^ Th17 normalised to the control group. **(C)** Naïve CD4^+^ T cells isolated from *Runx2^f/f^* or *Runx2^f/f^dLckCre^+^*mice cultured with neutrophils (0.5:1 neutrophil to CD4^+^ T cell ratio) in the presence or absence of Dnase1 for 72h under Th17 polarizing conditions. Representative IL-10 and GM-CSF in Th17 cells with analysis of the ratio of GM-CSF^+^ Th17 to IL-10^+^ Th17 cells in the presence or absence of Dnase1, normalising to control (*Runx2*^f/f^ Neutrophils) in each experiment. Symbols represents biological replicates. **(A)** Each symbol represents a biological replicate, data are mean ± SEM of n=9 mice per group from 2 independent experiments. Multiple unpaired t tests were used for comparisons. **(B)** Data are mean ± SEM of n=6 mice per group from two independent experiments and matched repeated measures one-way ANOVA with Dunnett corrections **(C)** Data are mean ± SEM of n=6 mice per group from two independent experiments. Two-way ANOVA with Sidak-Bonferroni corrections was utilized for comparisons. *P≤0.05, **P≤0.01 and *** P≤0.001.

## Discussion

Transcriptional regulation underlying pathogenic and stem-like Th17 cell states is a complex and incompletely understood process, which remains key to understanding their protective and pathogenic roles during homeostasis and tissue inflammation, respectively. Currently, the transcriptional control of pathogenic Th17 cells is better characterized compared to those that intrinsically suppress Th17 pathogenicity and/or maintain less inflammatory Th17 cell states ^10–12,37–39^. In this study, we identified the transcription factor RUNX2 is elevated in pathogenic CD4^+^ T cells in EAE and CSF CD4^+^ T cells during MS. RUNX2 was originally identified as a key osteoblast differentiation factor, and recently it has been studied in the context of various immune cells functions, including plasmacytoid dendritic cell egress, memory CD8^+^ T cell persistence and T-follicular helper cell differentiation ^21,23,40^. Its role in Th17 cells remains relatively unexplored, although it is notable that GM-CSF-secreting inflammatory CD4^+^ T cells in EAE have been previously determined to be enriched for open chromatin containing RUNX2 binding motifs ^41^. Among subsets of CD4^+^ T cells induced during EAE, we found RUNX2 was selectively enriched in GM-CSF-expressing cells, including pathogenic Th17 cells, in both the CNS and spleen. Its expression in Th17 cells can be attributed in part to regulation by JunB and BATF, which are well established Th17-specifying transcription factors ^42–44^. We demonstrate that *Runx2* deletion in T cells lowers the threshold of EAE induction and increases production of pathogenic Th17 cells in by integrating signals from cross-talk with neutrophils.

Overall, our findings indicate a novel role for RUNX2 as a negative regulator of Th17 pathogenicity. The observation that RUNX2 is expressed at low levels in non-pathogenic Th17 cells relative to their pathogenic counterparts in EAE and that its deletion in T cells increases pathogenic Th17 cells suggests that it functions as a cell-intrinsic brake in a negative feedback loop in Th17 cells to limit their pathogenicity. Similar negative feedback mechanisms in Th17 cells involving IL-24 and IL-17A, which limit Th17 pathogenicity through regulation of IL-10 and GM-CSF respectively, have previously been reported ^45,46^. However, specific transcriptional networks governing these cytokine mediated processes have not been elucidated, and whether RUNX2 is involved in the regulation of these processes or if it is part of a separate mechanism to regulate Th17 cell pathogenicity remains to be determined.

Our findings diverge somewhat from the findings of an earlier investigation by Jenkins and colleagues ^24^, which reported that *Runx2* deletion in activated TCR transgenic CD4^+^ T cells using retroviral-driven CRISPR gene deletion drove preferential commitment toward a Th17 lineage whilst inhibiting T-follicular helper differentiation *in vivo* ^24^. We did not find any role for RUNX2 in general Th17 specification either *in vitro* or *in vivo* using the EAE model and instead revealed a role for RUNX2 specifically in the skewing of Th17 cell responses to a pathogenic phenotype that is driven by interactions with neutrophils. Importantly, the study by Jenkins and colleagues ^24^ did not assess pathogenicity of Th17 cells. It is possible that the apparent divergence between these studies with respect to the role of RUNX2 in Th17 priming stems from the context and timing of *Runx2* deletion between these studies, so direct comparisons between that study and the present study present difficulties. Our study utilized a mouse strain whereby *Runx2* was deleted in post thymic mature T cells prior to their activation by TCR engagement and costimulatory signals, while Jenkins and colleagues ^24^ deleted *Runx2* post CD4^+^ T cell activation using retroviral transduction *in vitro* before subsequent sort transfer experiments. Therefore, our study assesses the *de novo* effect of RUNX2 deletion on naïve CD4^+^ T cell activation, differentiation, and commitment both *in vivo* and *in vitro*.

Importantly, the differences in clinical disease outcomes between the effect of RUNX2 deletion in the EAE models used in this study show that in settings in which pathogenic Th17 priming is limited, *Runx2* deficiency enables pathogenic Th17 cell generation at levels sufficient for disease induction. We show unequivocally that that this is due to a loss of RUNX2 modulated crosstalk between T cells and neutrophils that restrains Th17 cell pathogenicity. We identify extracellular DNA from neutrophils as a mediator of Th17 cell pathogenicity that is restrained by RUNX2. However, it should be noted that additional neutrophil-derived products may also contribute to this crosstalk. Previously, neutrophils have been shown to promote Th17 responses through production of IL-23 ^47^, CCL2, CCL20 ^48^, NOS ^17^, MHC-II, B7-1/2 ^49^, anti-microbial peptides ^15,16^, histones ^18^, and extracellular DNA ^36^. Whether these or other neutrophil-derived factors are the limiting factor in the LA-EAE model used in this study remains to be determined. Our data also indicate that immature neutrophil responses are boosted in the spleen by adjuvant stimulation during immune priming, and that these immature neutrophils selectively drive pathogenic Th17 cells. The relevance of these findings to pathogenic Th17 cell responses in humans is supported by recent clinical studies demonstrating a correlation between serum IL-17A and neutrophil degranulation products with new MS lesion development ^14^, and the emergence of the neutrophil-lymphocyte ratio (NLR) as a biomarker of MS severity ^50–52^ and a predictor of MS relapse risk ^53^. The NLR ratio is elevated during relapse compared to remission ^53^ and is further increased in progressive compared to relapsing-remitting MS ^51^. Therefore, while it remains to be definitively determined, there are multiple lines of evidence emerging that neutrophils contribute to inflammatory autoimmunity by influencing Th17 cell pathogenicity.

Our data supports a role for neutrophil-derived extracellular DNA in promoting pathogenic Th17 differentiation *in vitro* and this process was restrained by RUNX2 regulated processes in Th17 cells. These findings which indicate a pathogenic role for extracellular DNA in driving inflammatory Th17 responses are corroborated by recent findings which demonstrated that intraperitoneal Dnase1 administration from EAE onset protected mice against severe disease, and other findings which showed that Dnase1 treatment in a mouse model of psoriasis led to the attenuation of IL-17A production in the inflamed skin ^36,54^. Th17 cells have previously been reported to detect neutrophil extracellular traps through histone recognition on TLR2 ^18^. These studies showed a clear role for histones in triggering IL-17A production downstream of TLR2 signalling, the potential role of the extracellular DNA component of NETs in this process was not tested. The previous study by Wilson et al., ^18^ tested the role of histones/TLR2 in IL-17A induction of non-pathogenic Th17 cells, while the present study investigated the role of neutrophils in induction of GM-CSF in pathogenic Th17 cells. In our study of pathogenic Th17 cells, neutrophil-driven GM-CSF-expression by Th17 cells was not dependent on TLR2, which suggests that neutrophil-derived extracellular DNA must be detected by alternative danger recognition receptors, such as TLR4 ^54^, TLR9 ^55^, cGAS-STING ^56^ or the Ku complex ^57^. Indeed, TLR4 has previously been reported to drive encephalitogenic CD4^+^ T cell responses during EAE. In those studies, EAE-immunized *Rag1*-deficient mice reconstituted with TLR4-deficient CD4^+^ T cells displayed attenuated disease severity, and reduced Th17 responses in the CNS and peripheral lymphoid organs ^58,59^. TLR4 has previously been suggested to promote inflammatory Th17 cells via TLR4-RelA-miR-30a, however its exact ligands involved in promoting encephalitogenic T cell responses during EAE remains unclear ^18,60^. TLR4 has previously been implicated in extracellular DNA sensing by keratinocytes ^54^, in the sensing of chromatin associated nuclear protein HMGB1 by regulatory T cells ^61^, and in sensing of heat shock like protein HSP70L1 in dendritic cells ^62^. Considering that these proteins may be in complex with released extracellular DNA, it remains a possibility that Dnase1 treatment may affect their interactions with Th17 cells in co-cultures, therefore whether RUNX2 is involved in inhibiting the pathogenic Th17 program through sensing of these proteins remains to be investigated. Additionally, TLR9 has been shown to activate immune cells through the recognition of HMGB1 and RAGE in complex with DNA ^55^. However, previous work has demonstrated that TLR9 is protective in EAE, and TLR9 deficient mice displayed exacerbated EAE severity but no differences in Th17 cells at peripheral sites of priming ^63^. This suggests that TLR9 may not be involved in the neutrophil-driven pathogenic Th17 priming observed in the present study. Additionally, Wang and colleagues have recently demonstrated a DNA sensing mechanism by CD4^+^ T cells which involved the Ku complex, increased DNA sensing through this mechanism enhanced TCR-driven T cell activation and proliferation, resulting in their increased encephalitogenicity in the EAE model ^57^. In that study, Wang and colleagues demonstrated a redundant role for the cGAS-STING pathway in the sensing of DNA in CD4^+^ T cells ^57^, therefore whether RUNX2 may modulate neutrophil-driven Th17 pathogenicity through DNA-sensing via the Ku complex remains to be investigated. Collectively, the findings of our study implicate neutrophil-derived extracellular DNA as a possible ligand for neutrophil-driven Th17 pathogenicity during EAE, and support a role for RUNX2 regulated processes in Th17 cells in restraining their pathogenicity that is induced by neutrophil-derived extracellular DNA.

Importantly, inflammatory immune responses associated with both neutrophil and Th17 responses are not only implicated in the pathogenesis of CNS autoimmunity such as in MS ^14^, but are also implicated in other chronic inflammatory diseases such as arthritis ^64–67^, infectious colitis ^68–72^ and periodontitis ^73,74^, among others. Additionally, they are also implicated in protective immunity, in limiting the expansion of commensal microbes such as segmented filamentous bacteria (SFB) ^69,75^. The strong correlation between the extents of neutrophil-driven and Th17-driven inflammation in both pathological and protective contexts supports a model of bidirectional crosstalk whereby both cell types may positively or negatively regulate one another. Previous studies indicate that adaptive T cell production of IL-17A drives neutrophil mobilisation in response to microbial challenge ^75,76^. Additionally, Th17 cell supernatants have been shown to increase neutrophil expression of activation markers independent of IL-17A, and this was dependent on GM-CSF, TNF-α and IFN-γ ^48^. Conversely, as discussed previously there is strong *in vitro* evidence that neutrophils modulate Th17 polarization, and this is substantiated in a recent study demonstrating impaired peripheral Th17 responses in humans with genetic defects in neutrophil effector responses ^77^. Overall, neutrophil:Th17 crosstalk is vital in mediating protective and controlled inflammatory responses, however its dysregulation either through neutrophils or Th17 cells can lead to superfluous inflammatory responses underlying the development of chronic inflammatory diseases and autoimmunity. This study identifies *Runx2* as a regulator of pathways that limit Th17 pathogenicity that arises from neutrophil-Th17 crosstalk during EAE, the identification of therapeutics to enhance *Runx2*-driven pathways in peripheral Th17 cells to limit their pathogenicity may represent a potential intervention in MS and other forms of pathological inflammation driven by Th17 cells.

## Material and methods

### Study design

The role of the transcription factor RUNX2 in the regulation of pathogenic Th17 responses during autoimmune neuroinflammation was investigated. Complementary models of active EAE in conjunction with T cell specific knock-out mice were utilized to investigate the requirement of RUNX2 in EAE pathogenesis and pathogenic Th17 responses. Experiments involving the use of specific cell depletion antibodies and co-cultures of specific cell types were conducted to support cell-intrinsic roles. Data were analysed by flow cytometry and RNA-seq methods. No data points were excluded from analysis. The specific numbers of independent experiments, data normalization across independent experiments and statistical methods are detailed in the figure legends. The sample sizes for experiments were not pre-determined. Female mice aged between 7-13 weeks old were utilized for experiments and randomly assigned to experimental groups taking into consideration housing. When scoring clinical EAE, researchers were blinded to genotype and/or experimental group. Specific details of the reagents used in this study are provided in table S3. Study was conducted in accordance with ARRIVE Essential 10 guidelines.

### Mice

All mice were bred and housed under SPF conditions at the University of Adelaide Laboratory Animal Services (LAS). C57BL6/J mice were purchased from the Australian Research Council or Ozgene, *Il17a-gfp* mice (JAX 018472) ^78^ were kindly gifted by Professor Paul Foster (University of Newcastle, NSW), *dLck-hCre^3779^* mice ^27^ and *Runx2^f/f^* mice were provided by Professor Gabrielle T. Belz (University of Queensland). *Runx2^f/f^dLckCre^+^*mice were generated in house at LAS by intercrossing *Runx2^f/f^* and *dLck-hCre^3779^* mice. *Tlr2^-/-^* and control littermates ^79^ were bred and humanely euthanised at John Curtin School of Medical Research and splenocytes shipped to Adelaide on ice overnight for downstream experimentation. All mice were humanely euthanized by CO2 asphyxiation. Breeding and experimental procedures involving mice were carried out under The University of Adelaide institutional animal ethics approvals S-2015-225, S-2015-226, S-2019-058, and S-2021-098.

### Experimental autoimmune encephalomyelitis

EAE was induced in female mice 7-13 weeks of age by subcutaneous hind-flank immunizations with 100 µl of MOG_35-55_/CFA on day 0 and intra-peritoneal injection of pertussis toxin (300 ng / mouse) on days 0 and 2 post-immunization. For HA-EAE and LA-EAE, MOG_35-55_/CFA was prepared using CFA containing 8.33 mg/ml or 1 mg/ml desiccated *Mycobacterium tuberculosis* H37Rα (Voight Global Distribution), respectively, emulsified at a 1:1 ratio with MOG_35-55_ peptide (GL Biochem) diluted to 2 mg/ml in sterile PBS. Mice were scored on an EAE severity scale (0 = no symptoms, 1 = tail weakness, 2 = flaccid tail, 2.5 = hind-limb impairments, 3 = hind-limb paralysis, 4 = moribund). For neutrophil depletion EAE experiments, anti-Ly6G or isotype control antibodies were administered daily 50 µg per mouse via the intra-peritoneal route. Specific details of antibodies used are listed in Table S3.

### Preparation of single cell suspensions

Brains and spinal cords were digested in Collagenase D and Dnase I and mechanically dissociated using a gentleMACS Octo Dissociator (Miltenyi Biotec). Dissociated and digested tissue were filtered and washed through a 70 µm filter (Corning). Single-cell suspensions were then centrifuged at 1000xg in a 40% percoll gradient (Sigma) to separate out and remove remaining myelin debris. Spleens and inguinal lymph nodes were mashed through a 70 µm filter and erythrocytes lysed.

### Cell culture

Naïve CD4^+^ T cells were enriched from single-cell suspensions prepared from the spleens and inguinal lymph nodes of naïve mice using an EasySep™ Mouse Naïve CD4^+^ T Cell Isolation Kit (StemCell Technologies). Naïve CD4^+^ T cells were activated with plate bound anti-CD3 (5 µg/mL) and soluble anti-CD28 (1 µg/mL) and cultured for 3 days under specific polarizing cytokine conditions: Th17 (anti-IFNγ (10 µg/mL), anti-IL-4 (10 µg/mL), rm-IL-6 (25 ng/mL), rm-TGFβ1 (0.5 ng/mL), rm-IL-1β (25 ng/mL), rm-IL-23 (10 ng/mL) in glucose free RPMI media supplemented with D-(+)-Galactose (10 mM), FCS (10%), penicillin/streptomycin (1X), β-mercaptoethanol (0.27mM) (complete RPMI). For neutrophil-T cell co-cultures, neutrophils were enriched by negative selection from single-cell suspensions prepared from the spleens and inguinal lymph nodes of EAE-immunized mice using an EasySep™ Mouse Neutrophil Enrichment Kit (StemCell Technologies) as per manufacturer’s guidelines and a previous publication ^15^. Neutrophils were co-cultured at various ratios with naïve CD4^+^ T cells in the presence of plate bound anti-CD3, soluble anti-CD28 and Th17 polarizing cytokine conditions for 3 days. For experiments inhibiting extracellular DNA, Dnase1 (15U) was added to neutrophil:T cell co-cultures. Full details of antibodies and reagents are listed in Table S3.

### Flow cytometric analysis

For flow cytometric analysis of live splenocyte subsets, splenocytes were incubated with murine gamma globulin (Rockland) and viability dye (BD Horizon™ Fixable Viability Stain 780) prior to surface staining for CD45-BUV395, Ly6C-BV786, CD11b-PerCP-Cyanine5.5, TCRβ-BV421, IA/IE-BV510, Ly6G-FITC, CD19-PE-Dazzle, NK1.1-PE-Dazzle, CD11c-PE-Cy7, CD64-APC, CXCR2-APC, CD49d-PE. For flow cytometric analysis of CD4^+^ T cells, cells were restimulated with PMA (20 pg/ml), Ionomycin (1 nM), Golgistop (BD Biosciences) and Golgiplug (BD Biosciences) in IMDM supplemented with FCS (10%), Glutamax (1X), penicillin/streptomycin (1X), β-mercaptoethanol (0.27mM) (restimulation media) for 4 hours at 37°C. Cells were subsequently stained with viability dye (BD Horizon™ Fixable Viability Stain 780). For the detection of intracellular proteins, cytokines and transcription factors, cells were fixed and permeabilized using the Foxp3 / Transcription Factor Staining Buffer Set (eBioscience™) and stained with RUNX2-PE, CD45-BUV805, TCRβ-BUV395, CD4-BV786, CD44-BV480, IL-17A-BV421, IFNγ-FITC, GM-CSF-PE-Cy7, IL-10-PE-Dazzle and FOXP3-AF647. To enumerate cells, a fixed number of CountBright beads (Thermo Fisher) were acquired with each sample during sample acquisition. The ratio of beads to cells were calculated to determine absolute numbers of the cell populations of interest present in each sample. Flow cytometry data was acquired on a BD LSR Fortessa-X20 flow cytometer (BD Biosciences) and event acquisition was dependent on the beads gate as a stopping gate. All flow cytometry data was analysed with FlowJo software (Treestar). Gating strategies used for all analyses are detailed (Fig. S10-11). Full details of antibodies and reagents used for flow cytometry are listed in Table S3.

### ChIP-seq and ATAC-seq data and analysis

Th17 cell ATAC-seq and ChIP-seq datasets supporting the conclusions of this article are available in the ChIP-Atlas repository under the following accession codes: ATAC-seq (SRX15818878) ^12^, JunB ChIP-seq (SRX2773920) ^42^, BATF ChIP-seq (SRX187191) and Ep300 ChIP-seq (SRX187257) ^44^. Tracks were visualized on the Integrative Genomics Viewer (IGV) ^80^ against the *Runx2* gene on the MGSCv37 reference genome assembly.

### RNA-sequencing data and analysis

RNA-seq of CCR2^+^CCR6^-^ and CCR2^-^CCR6^+^ Th17 cells from EAE mice was carried out as follows. Briefly, CD4^+^ T cells were isolated from spleens of 3 female *Il17a-gfp* mice immunized for EAE and harvested on day 10 post-immunisation using an EasySep™ Mouse CD4 T cell Isolation Kit (StemCell Technologies) and subsequently viable cells were FACS-sorted using a BD FACSAriaIIImu cell sorter based on their expression of CD4, IL-17-GFP, CCR2 and/or CCR6. RNA was isolated from sorted cells using the Arcturus Pico Pure RNA Extraction Kit (Thermo Fisher) for bulk-RNA-sequencing using the the Illumina NextSeq platform. RNA sequencing of 150bp single end reads at an average of 15M reads per sample was performed. FASTQC ^81^ was utilized to ensure quality of raw reads and excess adaptor sequences were trimmed by cutadapt ^82^. Reads were subsequently aligned to GRCm38.93 by STAR Aligner ^83^ and counted by featureCounts ^84^, the resulting output of raw read counts were analyzed in R studio (v4.3.1). Differential gene expression analysis was carried out with the DESeq2 package (v1.12.3) ^85^. Other Th17 cell RNA-seq datasets utilized to support the conclusions of this manuscript are available in the Gene Expression Omnibus (GEO) repository under the following accession codes: GSE40918 ^44^, GSE98413 ^42^, GSE233908 ^13^ and GSE206304 ^12^.

### Statistical analysis

Data were analysed with GraphPad Prism 10.1.1, specific statistical tests, post-hoc tests, biological replicates and number of independent experiments are detailed in the specific figure legends. Briefly, paired t-tests, unpaired t-tests or ANOVA were used for parametric data, and Mann-Whitney U tests or Kruskal-Wallis H tests were used for non-parametric data. In all cases, p values <0.05 were considered statistically significant.

## Data and material availability

Data required to evaluate the conclusions of this manuscript are included in the paper and supplementary materials. Sequence data and raw read counts for RNA-seq of CCR2^+^CCR6^-^ and CCR2^-^CCR6^+^ Th17 cells that were generated in this study have been deposited in GEO with the primary accession code GSE296455. Requests for reagents, primary data and code should be directed to Dr Iain Comerford.

## Acknowledgements

We would like to thank Dr Ervin Kara (Rockefeller University, USA) for assistance with the Th17 cell RNAseq experiment; Dr Stevie Pederson (University of Adelaide) for assistance with preparatory analysis of RNAseq data; Professor Paul Foster (University of Newcastle) for providing *Il17a-gfp* mice; Professor Matthias Mack (University of Regensburg, Germany) for providing the anti-CCR2 antibody; staff at LAS (University of Adelaide) for assistance with animal husbandry; Dr Randall Grose (South Australian Health and Medical Research Institute) and Mrs Jessi Moore (University of Adelaide) for assistance with cell sorting. This study was supported by funds from MS Australia (21-3-012) and the NHMRC (APP1163335). IC was supported by senior research fellowship from MS Australia.

## Author Contributions

CWH performed experiments, analysed all data, prepared the figures and wrote the manuscript. AHH analysed RNAseq data and assisted. GTB and AB provided key reagents and essential advice. SRM co-supervised the study and wrote the manuscript. IC performed experiments, supervised the study and wrote the manuscript.

## Competing interests

The authors declare no competing interests.

## Notes

https://www.ncbi.nlm.nih.gov/geo/query/acc.cgi?acc=GSE296455

### Competing Interest Statement

The authors have declared no competing interest.

