## Supplemental Figures and Tables for "RUNX2 is a cell-intrinsic brake on Th17 pathogenicity driven by neutrophil extracellular traps"

**Supplementary Material**

**Supplemental Figures**

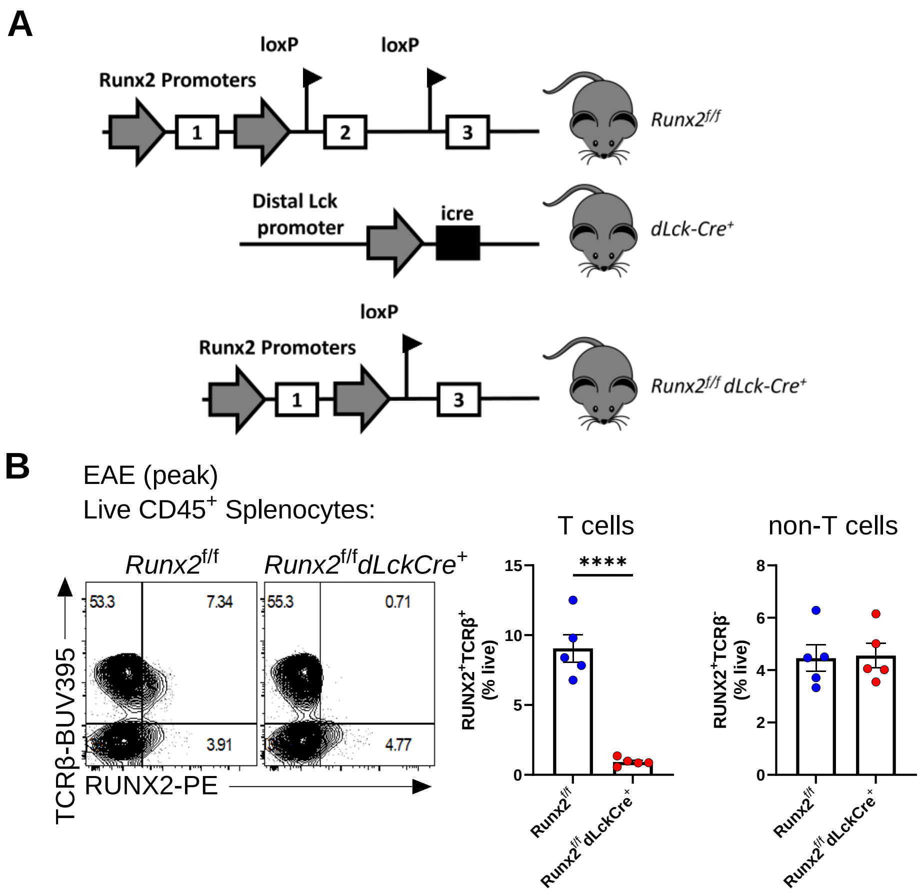


**Figure. S1. Validation of T cell conditional *Runx2*$^{-/-}$ mice.**

**(A)** Schematic of mice harbouring *Runx2* floxed alleles and mice with inducible Cre-recombinase under the control of the distal Lck promoter, and the resulting progeny when both are crossed. **(B)** Representative flow cytometry plots of TCRβ and RUNX2 co-expression on splenocytes from *Runx2*$^{f/f}$ and *Runx2*$^{f/f}$*dLckCre*$^{+}$ mice immunized for EAE (left) and quantification of the proportion of RUNX2$^{+}$TCRβ$^{+}$ and RUNX2$^{+}$TCRβ$^{-}$ cells (right). For cellular analysis each symbol represents a biological replicate, data are shown as mean ± SEM of n=5 mice per group across 2 independent experiments. Unpaired t-tests were used for comparisons. *P≤0.05, **P≤0.01 and *** P≤0.001

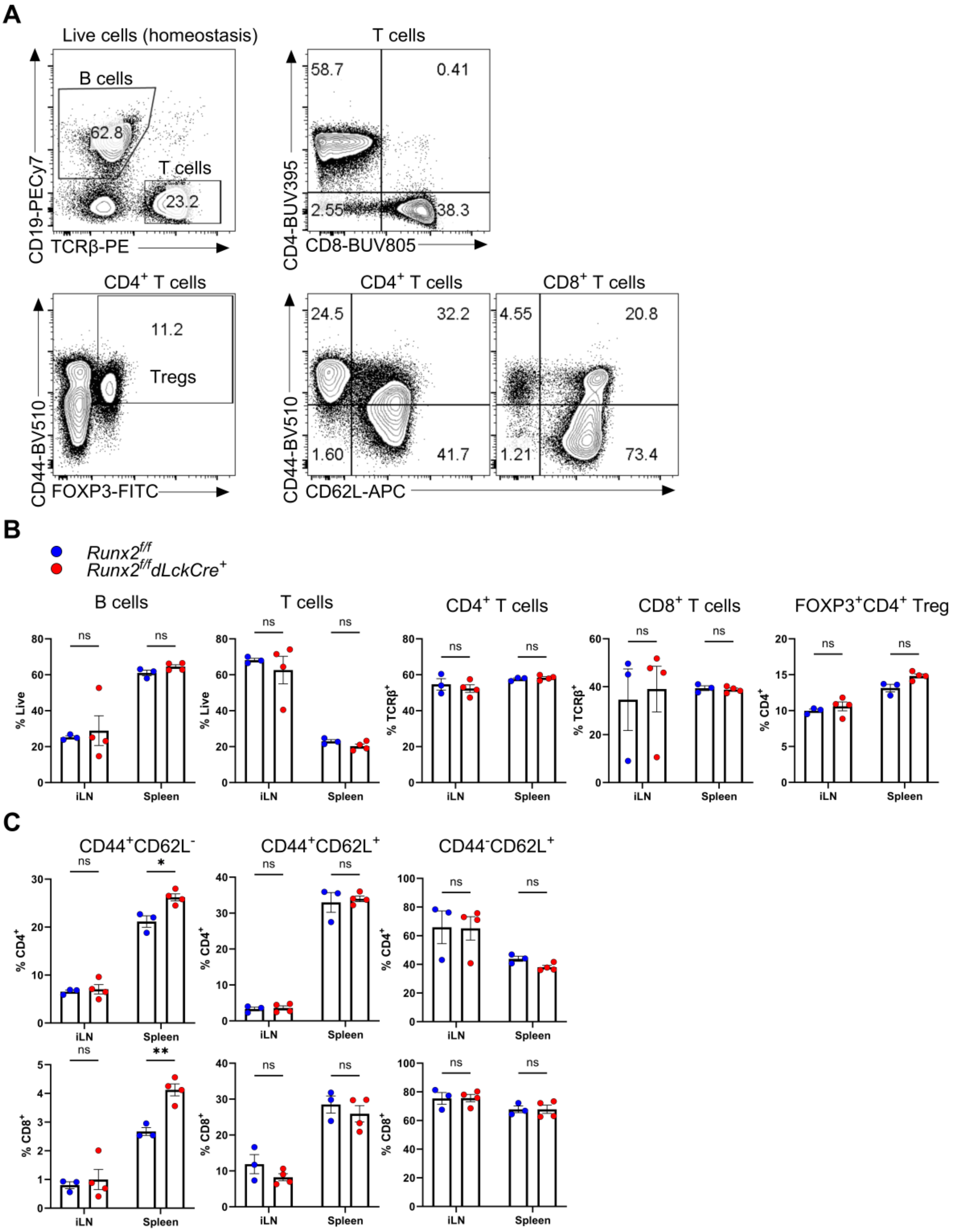


**Figure. S2. Analysis of lymphocytes at homeostasis in *Runx2$^{f/f}$dLckCre$^{+}$* mice.**

**(A)** Representative gating strategy used to determine CD19$^{+}$ (B cells), TCRβ$^{+}$ (T cells), CD4$^{+}$ T cells, CD8$^{+}$ T cells, CD4$^{+}$FOXP3$^{+}$ (Tregs), CD44$^{+}$CD62L$^{-}$ (effector memory), CD44$^{+}$CD62L$^{+}$ (central memory) and CD44$^{-}$CD62L$^{+}$ (naïve) CD4$^{+}$ and CD8$^{+}$ T cell populations in the spleen and inguinal lymph nodes at homeostasis in *Runx2$^{f/f}$ and Runx2$^{f/f}$dLckCre$^{+}$* mice. **(B)**

Quantification of B cells, T cells, $CD4^+$ T cells, $CD8^+$ T cells and Tregs. **(C)** Quantification of effector memory, central memory and naïve subsets in the $CD4^+$ and $CD8^+$ T cell compartment. For cellular analysis each symbol represents a biological replicate, data are shown as mean ± SEM of n=3 (*Runx2*$^{f/f}$) and n= 4 (*Runx2*$^{f/f}$*dLckCre*$^{+}$) mice across 2 independent experiments. Unpaired t-tests and Holm-Sidak post-hoc tests were used for comparisons. *P≤0.05, **P≤0.01 and *** P≤0.001

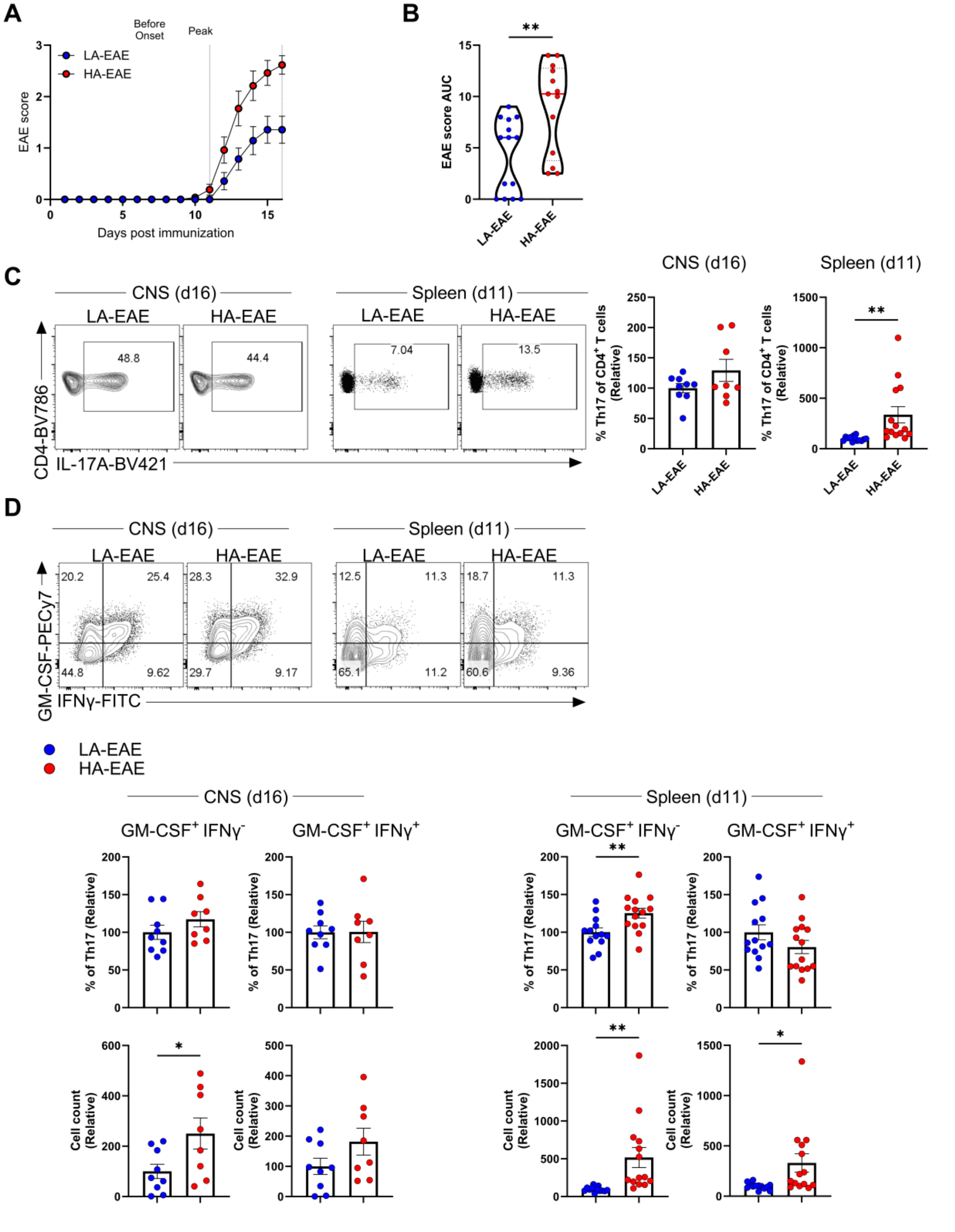


**Figure. S3. Adjuvant-induced modulation of pathogenic Th17 cell generation in EAE.** **(A)** EAE scores and **(B)** AUC analysis of wild-type mice immunized with either 50µg *M.tb* adjuvant/mouse (LA-EAE) or 415µg *M.tb* adjuvant/mouse (HA-EAE). EAE scores are shown as mean ± SEM of n=14 (LA-EAE) and n=13 (HA-EAE) across 3 independent experiments. Mann-Whitney tests were used for comparisons of AUC. **P≤0.01. **(C)** Representative flow cytometry plots of IL-17A expression on $CD4^+$ T cells from the CNS (peak disease, day 16) and spleen (pre-onset, day 11) of mice, the percentage of Th17 cells was calculated, normalising data to the LA-EAE group in each experiment. **(D)** Representative flow cytometry plots of GM-CSF and IFNγ expression on Th17 cells from the CNS (peak disease, day 16) and spleen (pre-onset, day 11) of mice, the percentage and number of $GM\text{-}CSF^+IFN\gamma^-$ and $GM\text{-}CSF^+IFN\gamma^+$ cells was calculated, defining cells in the LA group as 100% in each experiment. Each symbol represents a biological replicate, data of the CNS are shown as mean ± SEM of n=9 (LA-EAE) and n=8 (HA-EAE) across 2 independent experiments, data of the spleen are shown as mean ± SEM of n=13 (LA-EAE) and n=14 (HA-EAE) across 3 independent experiments. Unpaired t-tests were used for comparisons. *P≤0.05, **P≤0.01 and *** P≤0.001.

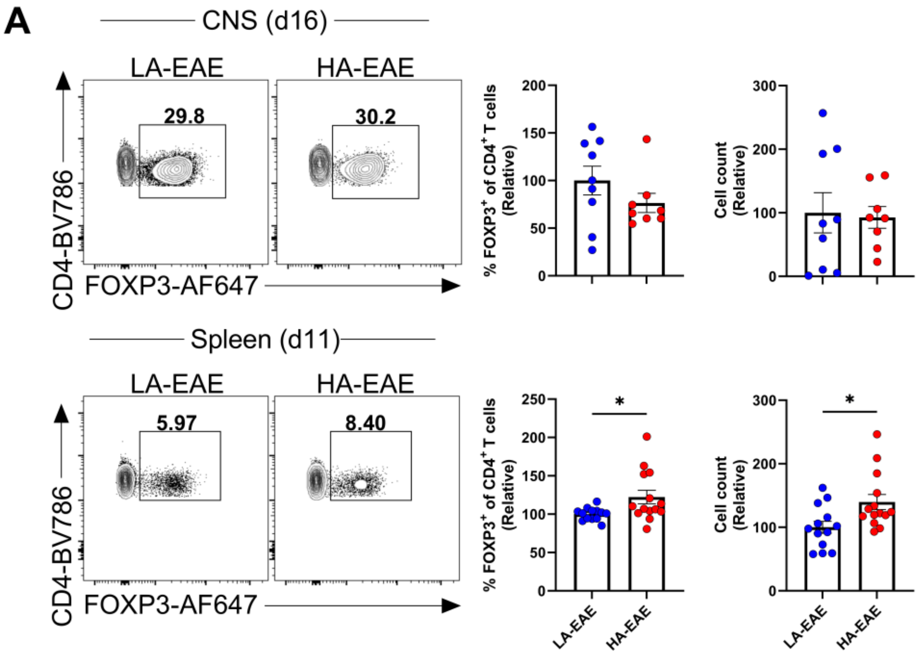


**Figure. S4. Regulatory T cell responses in LA-EAE and HA-EAE.**

**(A)** Representative flow cytometry plots and quantification of Treg cells in the CNS (peak EAE-day 16) and spleen (before EAE onset – day 11) of wild-type mice immunized for LA-EAE or HA-EAE. The percentage and number of FOXP3$^+$ cells were calculated, normalised to the LA-EAE group in each experiment. Each symbol represents a biological replicate, data of the CNS are shown as mean ± SEM of n=9 (LA-EAE) and n=8 (HA-EAE) across 2 independent experiments, data of the spleen are shown as mean ± SEM of n=13 (LA-EAE) and n=14 (HA-EAE) across 3 independent experiments. Unpaired t-tests were used for comparisons. *P≤0.05, **P≤0.01 and *** P≤0.001.

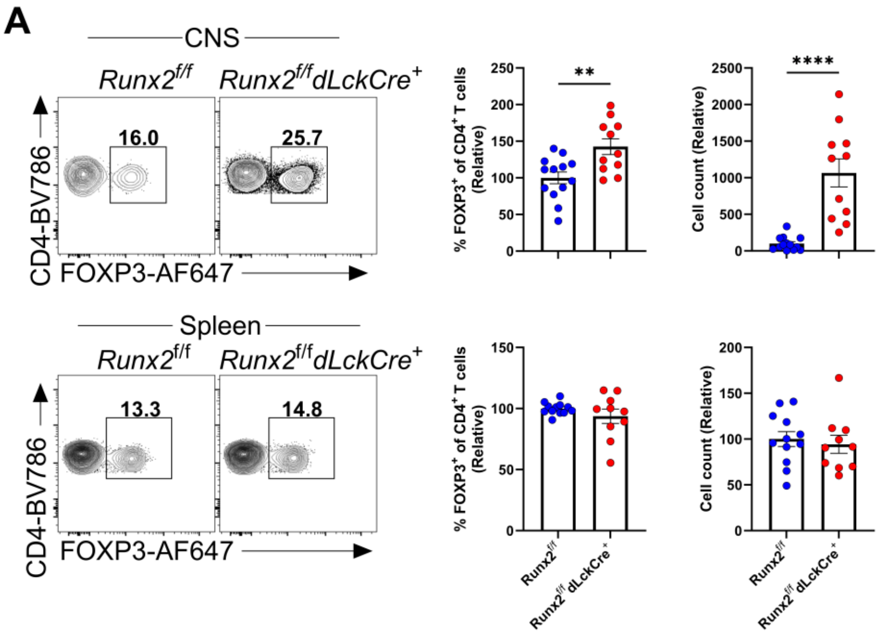


**Figure. S5. T cell deficiency of RUNX2 does not alter Treg generation in the spleen in LA- EAE.**

**(A**) Representative flow cytometry plots and quantification of Treg cells in the CNS and spleen of *Runx2*$^{f/f}$ and *Runx2*$^{f/f}$*dLckCre*$^{+}$ mice immunized for LA-EAE at day 13. The percentage and number of FOXP3$^{+}$ cells were calculated, normalised to the *Runx2*$^{f/f}$ group in each experiment. Each symbol represents a biological replicate, data are shown as mean ± SEM of n=12-13 (*Runx2*$^{f/f}$) and n=10-11 (*Runx2*$^{f/f}$*dLckCre*$^{+}$) mice across 4 independent experiments. Unpaired t-tests were used for comparisons. *P≤0.05, **P≤0.01 and *** P≤0.001.

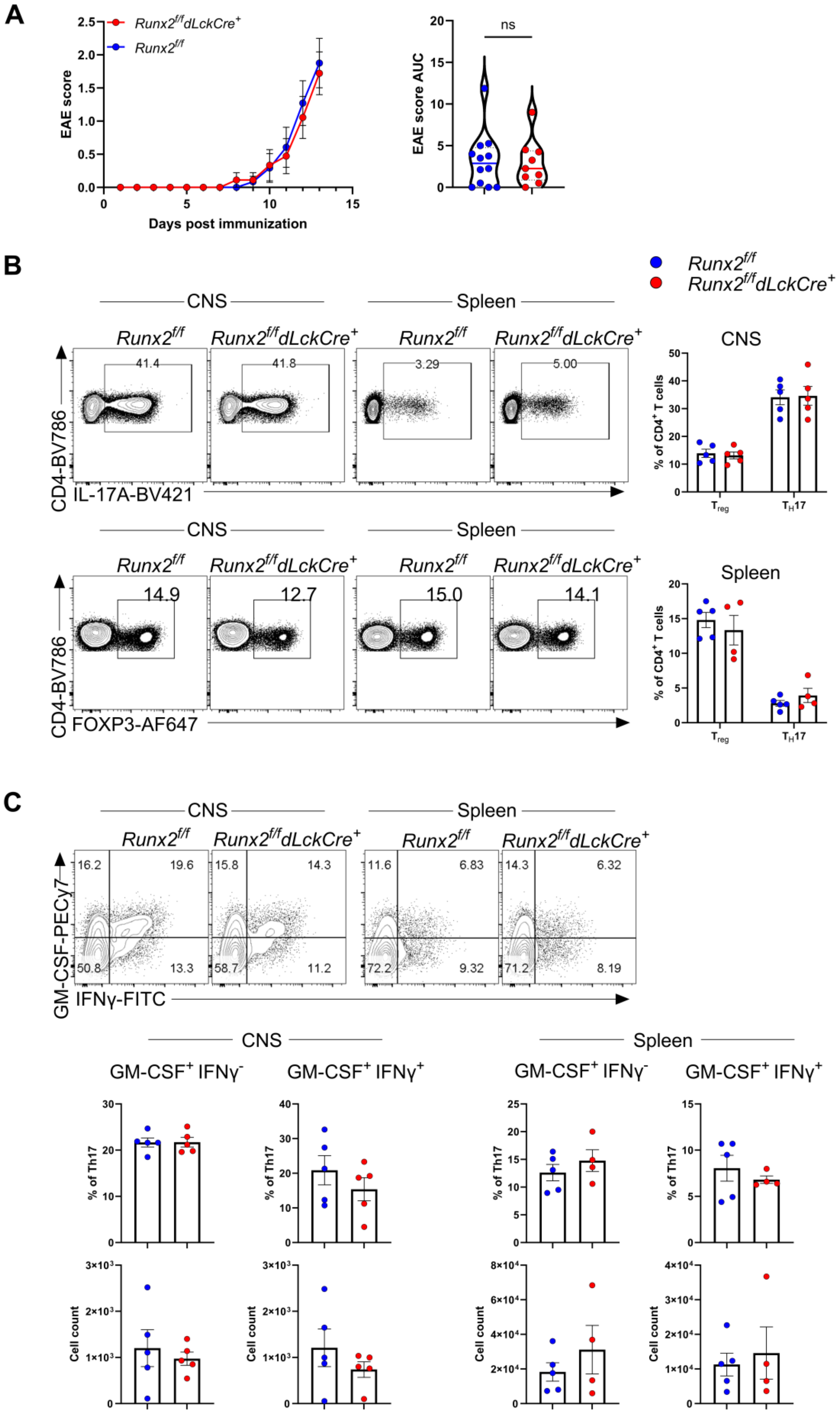
A
Runx2f/fdLckCre+
Runx2f/f
EAE score
Days post immunization
EAE score AUC
ns
B
Runx2f/f
Runx2f/fdLckCre+
CNS
Spleen
41.4
41.8
3.29
5.00
CD4-BV786
IL-17A-BV421
% of CD4+ T cells
Treg
TH17
14.9
12.7
15.0
14.1
FOXP3-AF647
C
16.2
19.6
15.8
14.3
11.6
6.83
14.3
6.32
50.8
13.3
58.7
11.2
72.2
9.32
71.2
8.19
GM-CSF-PECy7
IFNγ-FITC
GM-CSF+ IFNγ-
GM-CSF+ IFNγ+
% of Th17
Cell count

**Figure. S6. T cell deficiency of RUNX2 does not alter inflammatory and regulatory T cell responses in HA-EAE.**

**(A)** EAE scores and AUC analysis of *Runx2*$^{f/f}$ and *Runx2*$^{f/f}$*dLckCre*$^{+}$ mice immunized for HA-EAE. Each symbol represents a biological replicate, data are shown are mean ± SEM of n=12 (*Runx2*$^{f/f}$) and n=9 (*Runx2*$^{f/f}$*dLckCre*$^{+}$) across 3 independent experiments. **(B)** Representative flow cytometry plots of IL-17A and FOXP3 expression on CD4$^{+}$ T cells isolated from the CNS and spleen of *Runx2*$^{f/f}$ and *Runx2*$^{f/f}$*dLckCre*$^{+}$ mice immunized for HA-EAE at day 13 and percentages of Th17 and Treg cells. **(C)** Representative flow cytometry plots of GM-CSF and IFNγ expression in Th17 cells in the CNS and spleen and quantification of percentages and numbers. For cellular analysis each symbol represents a biological replicate, data are shown as mean ± SEM of n=4-5 mice per group from one experiment. Unpaired t-tests were used for comparisons.

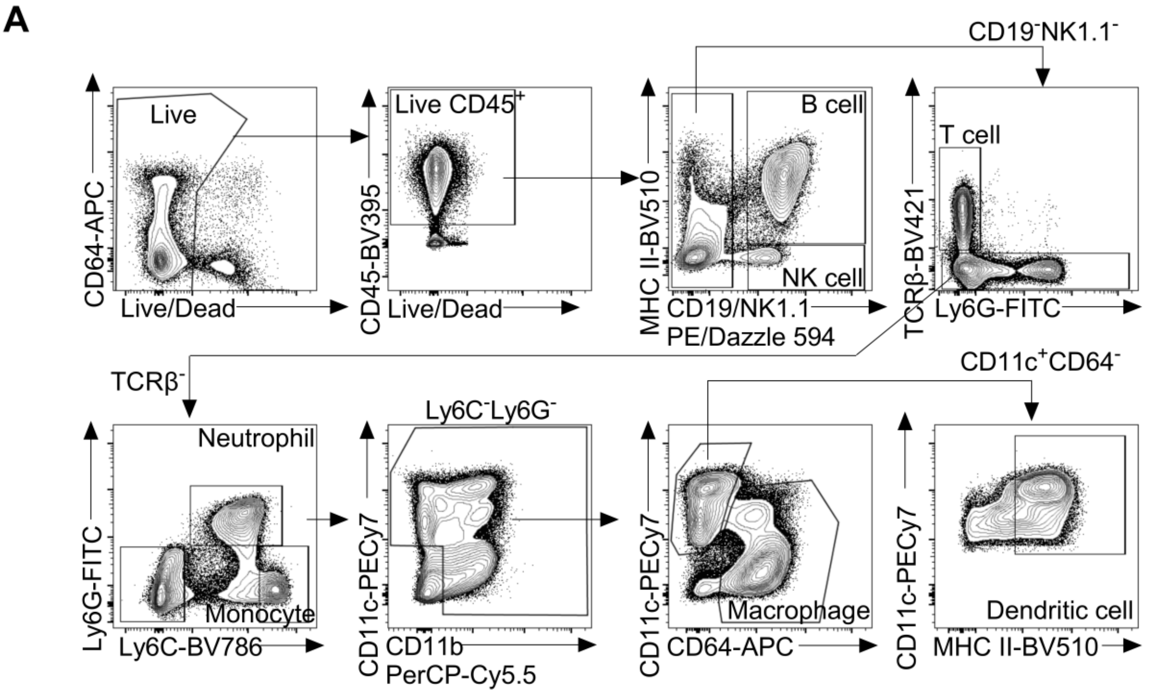


**Figure. S7. Gating strategy used to enumerate immune cell populations in the spleen.** **(A)** Representative flow cytometry gating strategy to identify B cells (CD45$^+$MHCII$^+$CD19$^+$), NK cells (CD45$^+$MHCII$^-$NK1.1$^+$), T cells (CD45$^+$CD19/NK1.1$^-$Ly6G$^-$TCRβ$^+$), neutrophils (CD45$^+$CD19/NK1.1$^-$TCRβ$^-$Ly6C$^{int}$Ly6G$^+$), Ly6C$^{hi}$ monocytes (CD45$^+$CD19/NK1.1$^-$TCRβ$^-$Ly6G$^-$Ly6C$^{hi}$), dendritic cells (CD45$^+$CD19/NK1.1$^-$TCRβ$^-$Ly6G$^-$Ly6C$^-$CD64$^-$CD11c$^+$MHCII$^+$) and macrophages (CD45$^+$CD19/NK1.1$^-$TCRβ$^-$Ly6G$^-$Ly6C$^-$CD11c$^+$CD11b$^+$CD64$^+$) in spleens of mice immunized for LA-EAE or HA-EAE and harvested on day 11

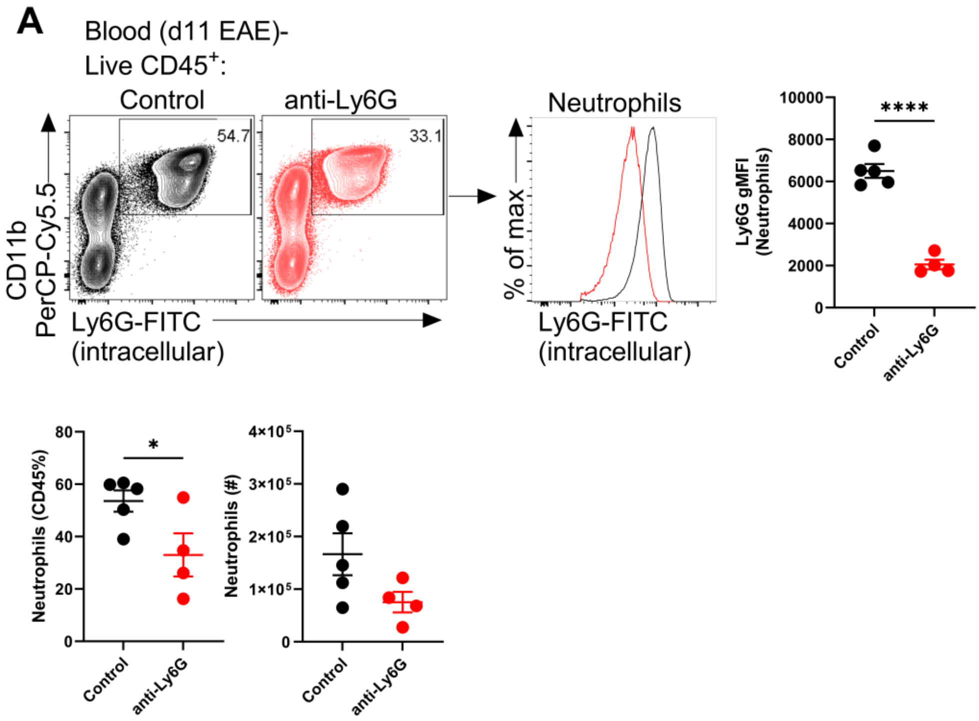


**Figure. S8. Depletion of neutrophils in EAE immunized mice.**

**(A)** Representative flow cytometry plots of CD11b and Ly6G expression on PBMCs isolated before EAE onset (day 11) from wild-type mice immunized for HA-EAE treated with either isotype control or anti-Ly6G antibodies daily. Quantification of the frequency and number of $CD11b^{+}Ly6G^{+}$ neutrophils and Ly6G geometric MFI on pre-gated neutrophils are shown. For cellular analysis each symbol represents a biological replicate, data are shown as mean ± SEM of n=5 (Isotype control) and n=4 (anti-Ly6G) treated mice from one experiment. Unpaired t-tests were used for comparisons. *P≤0.05, **P≤0.01 and *** P≤0.001.

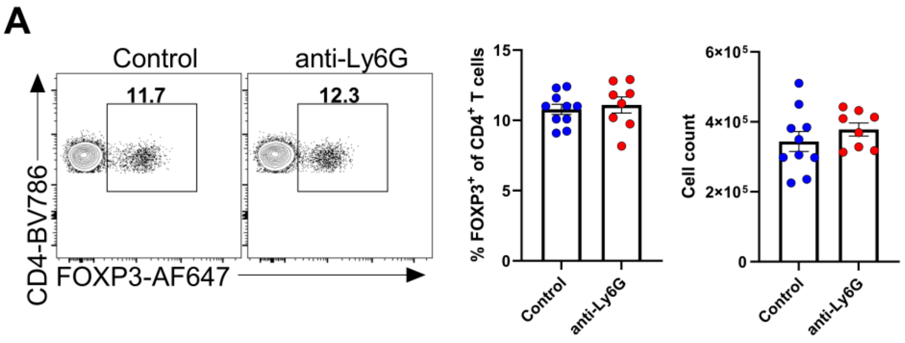


**Figure. S9. Neutrophil depletion does not alter regulatory T cells during EAE.**

**(A)** Representative flow cytometry plots of FOXP3 expression on $CD4^{+}$ T cells in the spleen and quantification of percentage and number of $FOXP3^{+}$ Treg cells. Each symbol represents a biological replicate, data are shown as mean ± SEM of n=8-10 mice per group across 2 independent experiments. Unpaired t-tests were used for comparisons. *P≤0.05, **P≤0.01 and *** P≤0.001.

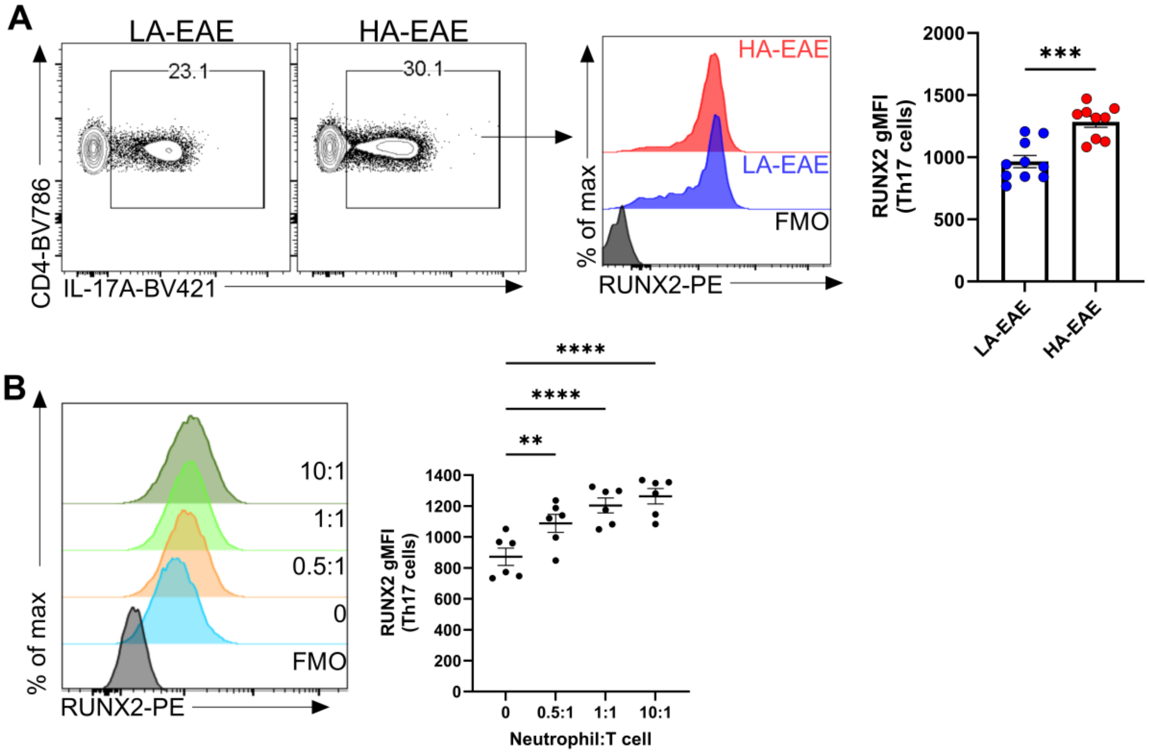


**Figure. S10. Neutrophils promote Th17 cell expression of RUNX2.**
**(A)** Representative histogram overlays of RUNX2 expression in Th17 cells from the spleens of mice immunized for LA-EAE and HA-EAE at day 11 and quantification of the geometric mean fluorescence intensity of RUNX2 in Th17 cells. **(B)** Representative flow cytometry plots of RUNX2 expression in Th17 cells co-cultured for 72h with neutrophils isolated from EAE mice. Quantification of the geometric mean fluorescence intensity of RUNX2 in Th17 cells. Each symbol represents a biological replicate. **(A)** Data are mean ± SEM of n=9-10 mice per group across two independent experiments and unpaired t-tests were used for comparisons. **(B)** Data are mean ± SEM of n=6 mice per group from two independent experiments and one-way ANOVA with Dunnett's corrections were used for comparisons. *P≤0.05, **P≤0.01 and *** P≤0.001.

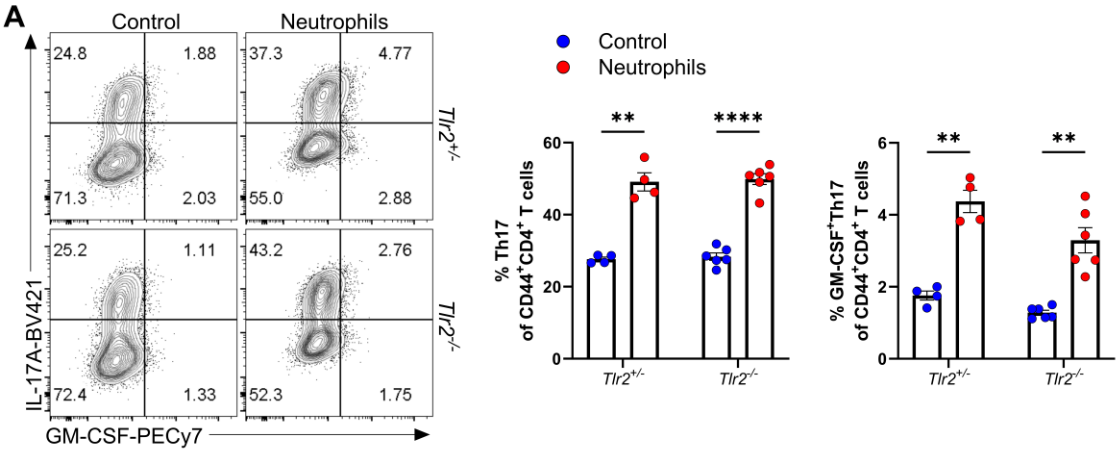


**Figure. S11. Neutrophil co-culture driven pathogenic Th17 responses are independent of TLR2-histone interactions.**

**(A)** Naïve $CD4^+$ T cells from *Tlr2*$^{+/-}$ or *Tlr2*$^{-/-}$ mice activated under Th17 polarizing conditions in the presence or absence of neutrophils. Representative plots of IL-17A and GM-CSF expression in $CD44^+CD4^+$ T cells. Quantification of Th17 and GM-CSF$^+$ Th17 frequencies. Each symbol represents a biological replicate. Data are mean ± SEM of n=4-6 mice per group from one experiment. Multiple paired t-tests with Holm Sidak corrections were used for statistical comparisons. *P≤0.05, **P≤0.01 and *** P≤0.001.

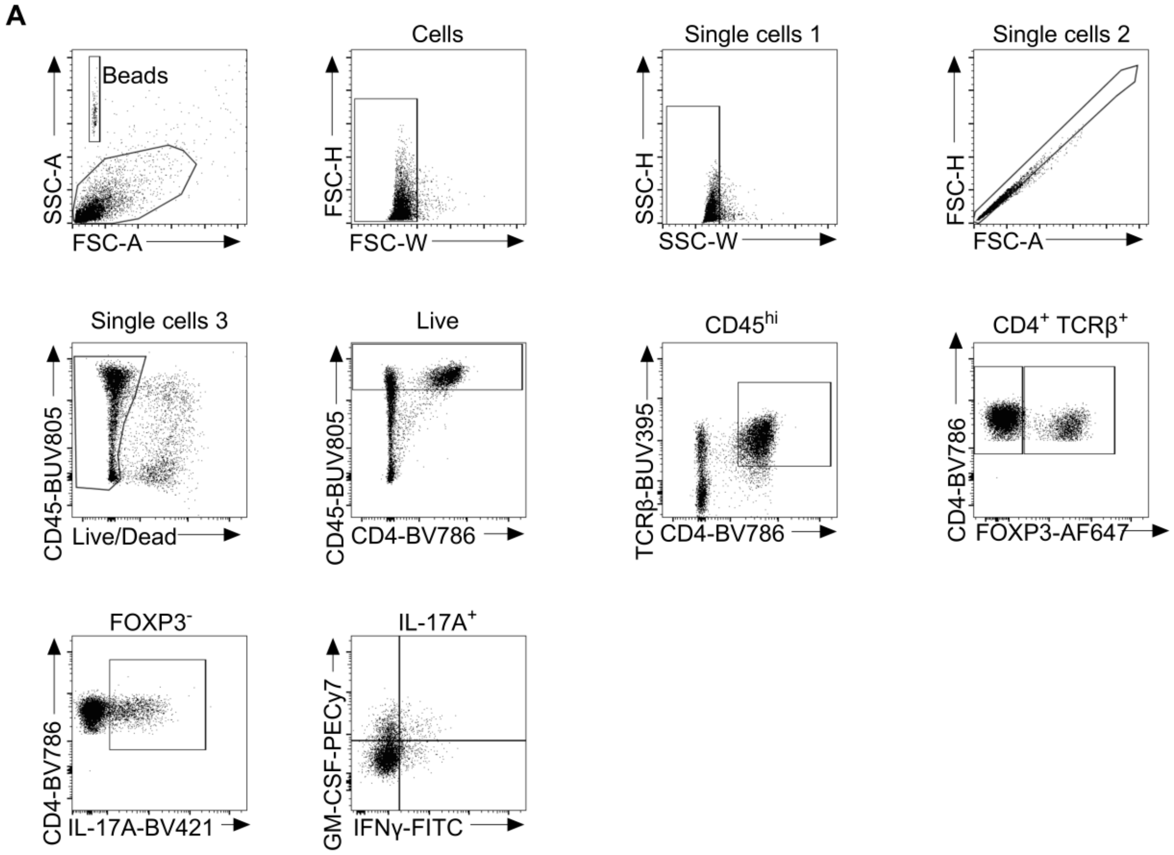


**Figure. S12. Gating strategies utilized for analysis of *in vivo* EAE experiments performed in this study.**

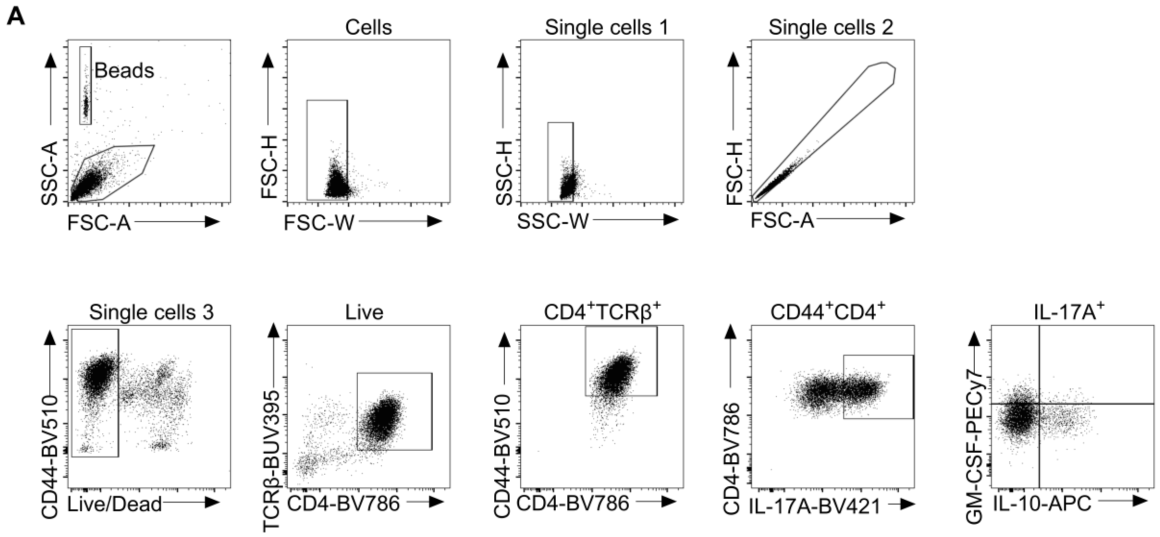


**Figure. S13. Gating strategy utilized for analysis of i*n vitro* T cell cultures performed in this study.**

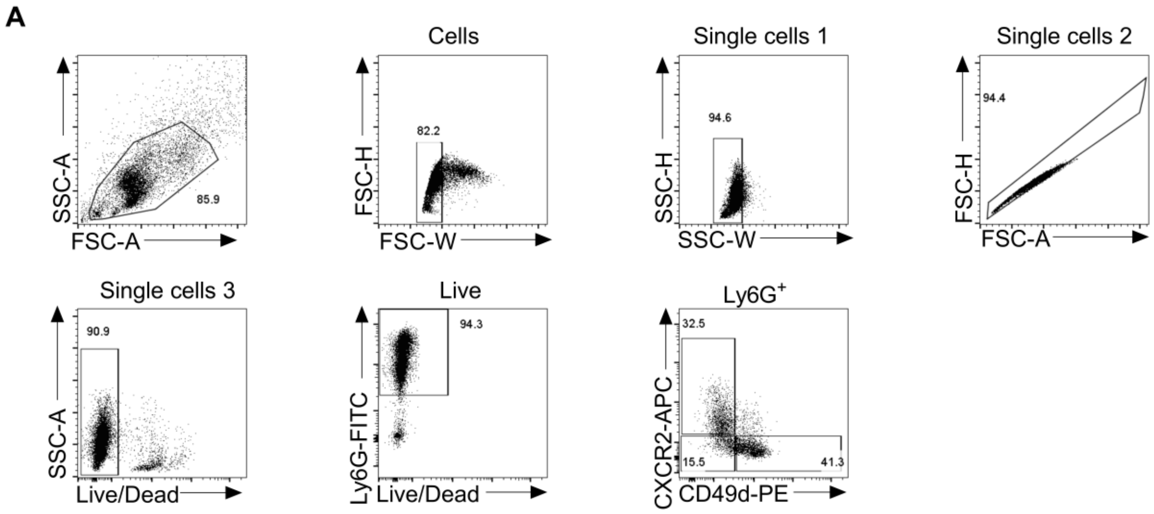


**Figure. S14. Gating strategies utilized for FACS-sorting of neutrophils in this study.**

**(A)** Neutrophils were enriched from the spleens of HA-EAE immunized mice at d11 post-immunization and FACS-sorted based on CXCR2 and/or CD49d expression.

**Supplemental Table**

**Table 1. Top upregulated and downregulated genes in $CCR2^{+}$ $CCR6^{-}$ compared to $CCR2^{-}$ $CCR6^{+}$ Th17 cells.**

| No | Gene name | Adjusted P value | No | Gene name | Adjusted P value |
|---|---|---|---|---|---|
| 1 | *Nkg7* | 1.78E-51 | 26 | *Hs3st3b1* | 1.89E-13 |
| 2 | *0610040F04Rik* | 1.43E-27 | 27 | *Lcn2* | 1.98E-13 |
| 3 | *Ifitm1* | 2.68E-27 | 28 | *Gcnt2* | 1.98E-13 |
| 4 | *Sema4a* | 6.78E-27 | 29 | *Ceacam1* | 4.11E-13 |
| 5 | *Ifng* | 2.27E-25 | 30 | *C5ar1* | 4.48E-13 |
| 6 | *Bhlhe40* | 2.17E-24 | 31 | *Cish* | 4.08E-12 |
| 7 | *Ctsw* | 9.30E-21 | 32 | *Epha8* | 5.89E-12 |
| 8 | *Tbx21* | 1.92E-20 | 33 | *Ermn* | 7.09E-12 |
| 9 | *Ifngr1* | 2.78E-19 | 34 | *Nupr1* | 4.24E-11 |
| 10 | *AA467197* | 6.14E-19 | 35 | *Chsy1* | 6.17E-11 |
| 11 | *Fcgr2b* | 3.21E-18 | 36 | *Hip1* | 1.20E-10 |
| 12 | *Ly6g5b* | 4.86E-18 | 37 | *Ccr2* | 3.21E-10 |
| 13 | *Lilrb4a* | 7.11E-18 | 38 | *Tasl* | 3.28E-10 |
| 14 | *Trim16* | 2.15E-17 | 39 | *Anxa1* | 5.30E-10 |
| 15 | *Il18rap* | 2.62E-16 | 40 | *Il12rb2* | 1.01E-09 |
| 16 | *Tns2* | 6.39E-16 | 41 | *Hid1* | 1.29E-09 |
| 17 | *Swap70* | 1.09E-15 | 42 | *Il2ra* | 5.44E-09 |
| 18 | *Runx2* | 1.15E-15 | 43 | *Sytl2* | 9.34E-09 |
| 19 | *Cxcr6* | 1.58E-14 | 44 | *Fes* | 9.51E-09 |
| 20 | *Tbc1d2* | 2.34E-14 | 45 | *Elane* | 1.05E-08 |
| 21 | *Ltf* | 2.42E-14 | 46 | *Lgals3* | 1.16E-08 |
| 22 | *Cd177* | 5.66E-14 | 47 | *Tm6sf1* | 1.51E-08 |
| 23 | *Itgb2l* | 7.27E-14 | 48 | *Coro2a* | 2.01E-08 |
| 24 | *Gzmb* | 1.01E-13 | 49 | *Csf1* | 2.61E-08 |
| 25 | *Chil1* | 1.18E-13 | 50 | *Wdr95* | 4.74E-08 |

| No | Gene name | Adjusted P value | No | Gene name | Adjusted P value |
|---|---|---|---|---|---|
| 1 | *Tox2* | 1.58E-12 | 26 | *Nme4* | 8.00E-05 |
| 2 | *Il6st* | 1.04E-11 | 27 | *Ube2e2* | 8.43E-05 |
| 3 | *Btla* | 1.33E-11 | 28 | *Rflnb* | 0.000106 |
| 4 | *Tnfsf8* | 1.78E-11 | 29 | *Ldhb* | 0.000107 |
| 5 | *Adk* | 1.92E-10 | 30 | *Asb2* | 0.000107 |
| 6 | *Thada* | 5.39E-10 | 31 | *Asap1* | 0.000124 |
| 7 | *Fcrl1* | 1.16E-08 | 32 | *Actn1* | 0.000152 |
| 8 | *Nipal1* | 1.16E-08 | 33 | *BC106179* | 0.000152 |
| 9 | *Nav2* | 1.77E-08 | 34 | *Slc16a5* | 0.000185 |
| 10 | *Ccr6* | 4.21E-08 | 35 | *Ddr1* | 0.000217 |
| 11 | *Aff3* | 1.07E-07 | 36 | *Afp* | 0.000243 |
| 12 | *Bcl2* | 2.95E-07 | 37 | *Ifi27l2a* | 0.000245 |
| 13 | *Il23r* | 1.29E-06 | 38 | *Scarb1* | 0.000251 |
| 14 | *Art2b* | 1.32E-06 | 39 | *Nrgn* | 0.000313 |
| 15 | *Il22b* | 2.69E-06 | 40 | *Patj* | 0.000351 |
| 16 | *Plekha1* | 4.30E-06 | 41 | *Hpgds* | 0.000356 |
| 17 | *Ramp3* | 7.09E-06 | 42 | *Phactr2* | 0.000369 |
| 18 | *A630023P12Rik* | 8.99E-06 | 43 | *Snn* | 0.000405 |
| 19 | *St6gal1* | 1.42E-05 | 44 | *H2-Ob* | 0.000442 |
| 20 | *Gm11454* | 2.60E-05 | 45 | *Scml4* | 0.000523 |
| 21 | *Il9r* | 2.96E-05 | 46 | *Gm14718* | 0.000531 |
| 22 | *Pou2af1* | 3.65E-05 | 47 | *Mgst2* | 0.000587 |
| 23 | *Tox* | 4.01E-05 | 48 | *Rgs10* | 0.000613 |
| 24 | *Cnn3* | 6.07E-05 | 49 | *Slamf6* | 0.000623 |
| 25 | *Trbv15* | 6.79E-05 | 50 | *Irf6* | 0.000648 |

$CCR2^+CCR6^-$ and $CCR2^-CCR6^+$ Th17 cells were assessed utilizing DESeq2 to determine top 50 differentially expressed genes in either direction. Top upregulated and downregulated genes were defined based on log2foldchange ≥ ±1.5 and adjusted p-value < 0.05.

**Table 2. Clinical disease parameters of *Runx2$^{f/f}$* and *Runx2$^{f/f}$dLckCre$^{+}$* mice immunized for LA-EAE or HA-EAE.**

| Genotype | Adjuvant | N (total) | Incidence (%) | Day of Onset | Max. Score |
|---|---|---|---|---|---|
| *Runx2$^{f/f}$* | LA | 20 | 60 | 15.45 ± 1.959 | 1.417 ± 0.6686 |
| *Runx2$^{f/f}$dLckCre$^{+}$* | LA | 22 | 86 | 13.91 ± 2.136 | 2.158 ± 0.7275 |
| Result of statistical test | | | * | * | ** |
| *Runx2$^{f/f}$* | HA | 12 | 75 | 11.44 ± 1.333 | 1.875 ± 1.295 |
| *Runx2$^{f/f}$dLckCre$^{+}$* | HA | 9 | 89 | 11.38 ± 1.685 | 1.722 ± 0.9718 |
| Result of statistical test | | | ns | ns | ns |

EAE incidence (%), average day of EAE onset ± SD and average maximal EAE score ± SD are reported. Statistical analysis (log-rank test comparing EAE incidence over time; Mann-Whitney tests comparing day of EAE onset and maximal EAE scores).

**Table 3. Antibodies and reagents used in this study.**

| Antibodies | Source | Clone; Catalog # |
|---|---|---|
| αCCR2-Biotinylated | Provided by Prof. Matthius Mack, Regensburg, Germany [86]. Biotinylated in house. | MC21 |
| αCCR6-PE | Biolegend | 29-2L17; 129803 |
| αCD45-BUV395 | BD Biosciences | 30-F11; 564279 |
| αLy6C-BV786 | BD Biosciences | HK1.4; 755197 |
| αCD11b-PerCP-Cyanine5.5 | Biolegend | M1/70; 101228 |
| αTCRβ-BV421 | Biolegend | H57-597; 109230 |
| αIA/IE-BV510 | BD Biosciences | M5/114.15.2; 742893 |
| αLy6G-FITC | BD Biosciences | 1A8; 11-9668-82 |
| αCD19-PE-Dazzle | Biolegend | 6D5; 115553 |
| αNK1.1-PE-Dazzle | BD Biosciences | PK136; 562864 |
| αCD11c-PE-Cy7 | Biolegend | N418; 117318 |
| αCD64-APC | Biolegend | X54-5/7.1; 139306 |
| αCXCR2-APC | Biolegend | SA044G4; 149312 |
| αCD49d-PE | Biolegend | 9C10(MFR4.B); 103706 |
| αRUNX2-PE | Cell Signalling | D1L7F; 98059 |
| αCD45-BUV805 | BD Biosciences | HM48-1; 568336 |
| αTCRβ-BUV395 | BD Biosciences | H57-597; 569248 |
| αCD4-BV786 | BD Biosciences | RM4-5; 563727 |
| αCD44-BV480 | BD Biosciences | IM7; 566200 |
| αIL-17A-BV421 | Biolegend | TC11-18H10.1; 506925 |
| αIFNγ-FITC | BD Biosciences | XMG1.2; 554411 |
| αGM-CSF-PE-Cy7 | Biolegend | MP1-22E9; 505412 |
| αIL-10-PE-Dazzle | Biolegend | JES5-16E3; 505034 |
| αFOXP3-AF647 | Biolegend | MF-14; 126408 |

| Reagents | Company | Clone; Catalog # |
|---|---|---|
| BD Horizon™ Fixable Viability Stain 780 | BD Biosciences | 565388 |
| eBioscience™ Foxp3 / Transcription Factor Staining Buffer Set | Invitrogen | 00-5523-00 |
| EasySep™ Mouse CD4+ T Cell Isolation Kit | Stemcell | 19852 |
| EasySep™ Mouse Naïve CD4+ T Cell Isolation Kit | Stemcell | 19765 |
| EasySep™ Mouse Neutrophil Enrichment Kit | Stemcell | 19762 |
| InVivoMAb anti-mouse CD3 | BioXcell | 17A2; BE0002 |
| BD Pharmingen™ Purified Hamster Anti-Mouse CD28 | BD Biosciences | 37.51; 553295 |
| InVivoPlus anti-mouse IFNγ | BioXcell | XMG1.2; BP0055 |
| InVivoMAb anti-mouse IL-4 | BioXcell | 11B11; BE0045 |
| rm-IL-6 | R&D Systems | 406-ML |
| rm-TGFβ1 | R&D Systems | 7666-MB |
| rm-IL-1β | Biolegend | 575106 |
| rm-IL-23 | Biolegend | 589006 |
| Pertussis toxin | Sapphire | 181 |
| $MOG_{35-55}$ | GL Biochem (Shanghai) Ltd | 51716 |
| Complete Freund's adjuvant | Sigma-Aldrich | F5881-6X10ML |
| Desiccated Mycobacterium tuberculosis H37Rα | BD Biosciences | Difco-231141 |
| Bioxcell InVivoMAb anti-mouse Ly6G | BioXcell | 1A8; BE0075-1 |
| Bioxcell InVivoMAb rat IgG2a isotype control, anti-trinitrophenol 25MG | BioXcell | BE0089 |
| Collagenase D | Sigma | 11088858001 |
| DNAse I | Sigma | D5025-150KU |
| PMA | Sigma | P1585-1MG |

| | | |
|---|---|---|
| Ionomycin calcium salt from Streptomyces conglobatus | Sigma | I0634-1MG |
| Golgi stop | BD | 554724 |
| Golgi plug | BD | 555029 |
| Arcturus Pico Pure RNA Extraction kit | Applied Biosystems/Thermo Fisher Scientific, Carlsbad, CA, USA | KIT0204 |
| Glucose free RPMI | Thermo | 11879020 |
| IMDM powder | Thermo | 12200036 |
| D-(+)-Galactose | Sigma | G0750-10G |
| Penicillin/Streptomycin | Thermo | 15140122 |
| Glutamax | Thermo | 35050061 |
| 2-mercaptoethanol | Sigma | M6250 |
| FCS | Sigma | 12003C |

296
